# Distinct mechanistic pathways of early tauopathy revealed by *MAPT* mutations

**DOI:** 10.64898/2026.02.14.705716

**Authors:** M.S. Foiani, R.S. Nirujogi, N. Watamura, S. Bez, E. Tsefou, A. Santambrogio, S. Patel, H. Davies, C. Goulbourne, N. Fatima, T. Birkle, E. Camporesi, G. Brinkmalm, H. Zetterberg, M. Wilkinson, S.E. Radford, N.A. Ranson, P. Maglio-Cauhy, E. Turkes, A. Avdic-Belltheus, I. Rawlinson, X. Prebibaj, L. Panti, D. Gavriouchkina, M. Blunskyte-Hendley, T. Saido, M. Vendruscolo, R. Frank, M. Bourdenx, K.E. Duff

## Abstract

Tau pathology underlies a broad spectrum of neurodegenerative disorders, collectively termed tauopathies, yet these diseases exhibit striking heterogeneity in their biological mechanisms and clinical outcomes. The basis for this heterogeneity remains poorly understood. Here, we address this question using knock-in mouse models expressing two distinct frontotemporal dementia-associated tau mutations to define how different tau variants drive divergent pathogenic programs *in vivo*. We find that the two mutations give rise to fundamentally different trajectories of tau pathogenesis. One trajectory is marked by progressive tau hyperphosphorylation and cytoskeletal destabilization occurring in the absence of detectable tau seed formation. In contrast, an alternative trajectory is characterized by tau hypophosphorylation, early seed formation, and alterations in nucleotide metabolism and chromatin organization, without overt cytoskeletal disruption. With aging, tau in this latter pathway transitions to a hyperphosphorylated state and forms mature fibrillar aggregates. Genetic enhancement of β-amyloid selectively accelerates fibril formation, particularly in the model exhibiting early seeding. Together, these findings demonstrate that distinct tau mutations can engage separable pathogenic mechanisms, providing a biological framework for the heterogeneity observed across tauopathies, and highlighting the need for mechanism-informed therapeutic strategies and patient stratification.

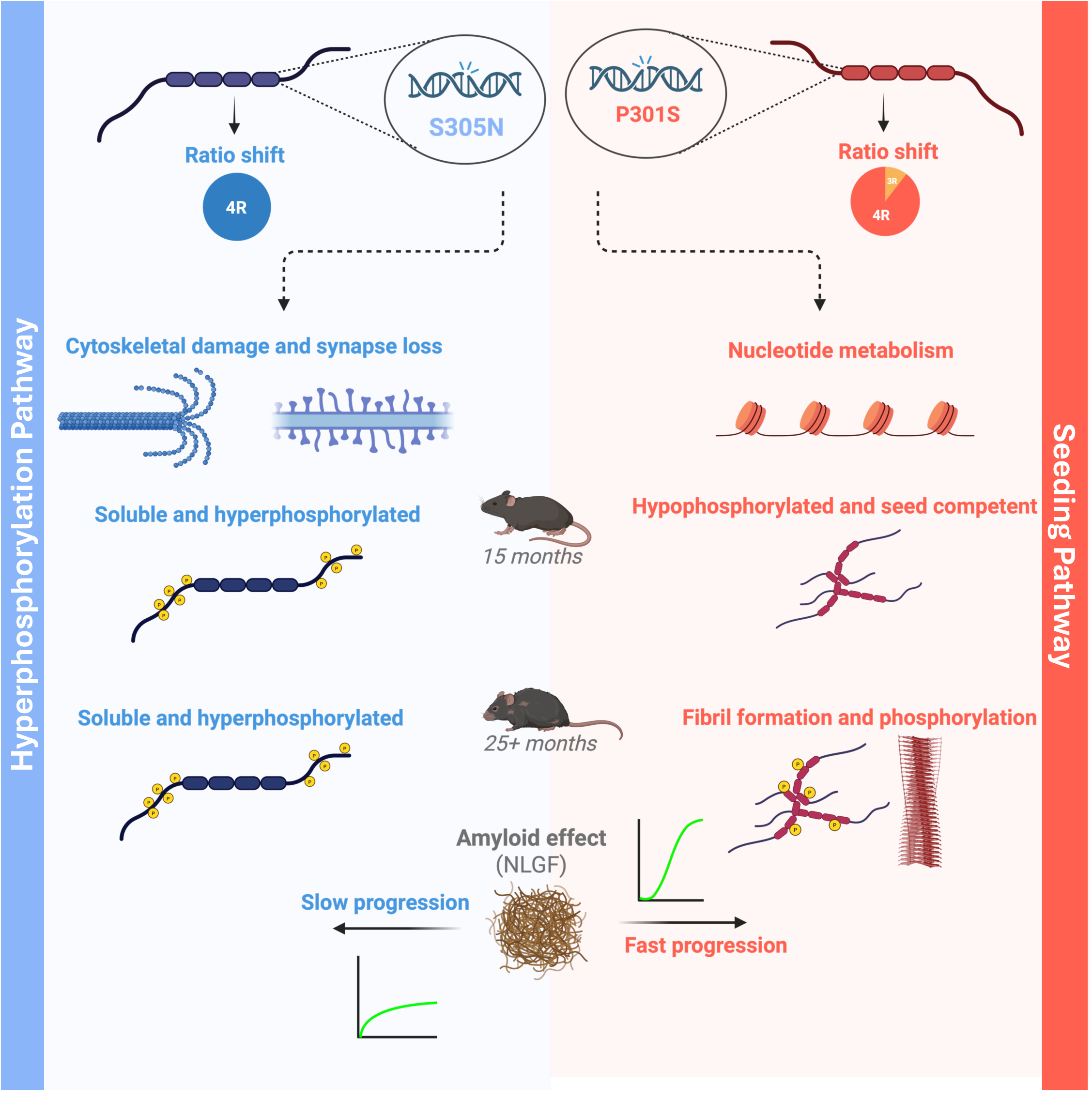

**Graphical abstract:** Schematic representation comparing two trajectories of tau pathology. The S305N mutation promotes a 4R isoform shift, cytoskeletal damage and synapse loss, and accumulation of soluble hyperphosphorylated tau. Tau remains soluble even at old ages. In contrast, the P301S mutation generates hypophosphorylated, seed-competent tau that forms fibrils. The effect of amyloid is slow in the S305N, but results in accelerated acceleration of pathology in the P301S. Figure made with Biorender.com.

## Introduction

Tauopathies are neurodegenerative diseases defined by intracellular accumulation of the microtubule-associated protein tau (*MAPT*). They encompass diverse clinically and pathologically distinct conditions, including Alzheimer’s disease (AD) and the frontotemporal lobar degeneration (FTLD) spectrum, the latter comprising progressive supranuclear palsy, corticobasal degeneration, Pick’s disease, and globular glial tauopathy. Alternative splicing of *MAPT* exon 10 yields tau isoforms with three or four microtubule-binding repeats (3R/4R)^1^ and any shift in the usual ratio of 1:1 3R:4R can lead to tauopathy. Tau pathology is characterized by the mislocalization of soluble tau from axons to somatodendritic compartments, where it accumulates as hyperphosphorylated, insoluble fibrils in neurons and glia. Pathogenic tau accumulation correlates with neurodegeneration, synapse loss, and depending on the region impacted, behavior, language, motor or cognitive deficits. The mechanisms that initiate the pathological cascade remain poorly understood, in part due to a scarcity of physiologically relevant models capturing the earliest events^2,3^.

Mutations in the tau gene (*MAPT*) are a major cause of familial FTLD and they provide insight into the mechanisms driving tau-mediated neurotoxicity that are likely to also be occurring in sporadic cases. Despite a common tau pathology at *post mortem*, both *MAPT* mutation carriers, and those with sporadic disease show striking phenotypic heterogeneity^4^. To date, more than 70 *MAPT* mutations have been reported^5^, altering isoform balance, structure, post-translational modification, and aggregation, which track with clinical symptoms^6,7^, biomarker profiles^8–11^, disease progression rates^12^, and severity^13^. Dissecting the mechanistic basis of this heterogeneity is essential for diagnosis, patient stratification, and targeted therapy.

Many tau transgenic mouse lines have been created to model tauopathy^14–19^, but overexpression introduces confounds, including supraphysiologic tau levels leading to rapid, overt pathology development, imbalanced isoforms, insertional effects, and challenges in generating matched controls^20^. Knock-in (KI) strategies that target the endogenous Mapt locus^21,22^, especially those that substitute a full-length human version of tau^23–25^, better preserve native spatial and temporal regulation and isoform expression.

To investigate the molecular causes of clinical and biological heterogeneity in tauopathies, we used two *MAPT* KI mouse lines harboring distinct mutations, *MAPT* S305N and P301S (**Fig. 1A**). Using this approach, we demonstrate that different *MAPT* mutations selectively disrupt distinct cellular functions, indicating mutation-specific biological consequences, rather than converging on a single pathogenic pathway. These differences were accompanied by the emergence of distinct tau profiles; in S305N mice, phosphorylation was sufficient to cause extensive cytoskeletal damage without the presence of aggregates. In P301S mice, non-phosphorylated seed-competent tau is the culprit, affecting nucleotide metabolism and chromatin organization. Finally, crossing *MAPT* KI mice to amyloid precursor protein (*APP*) KI mice with elevated β-amyloid, selectively accelerated seed and fibril formation. These results argue for mechanism-specific trajectories of tau pathogenesis with implications for target-selection and patient stratification.

**Figure 1.**
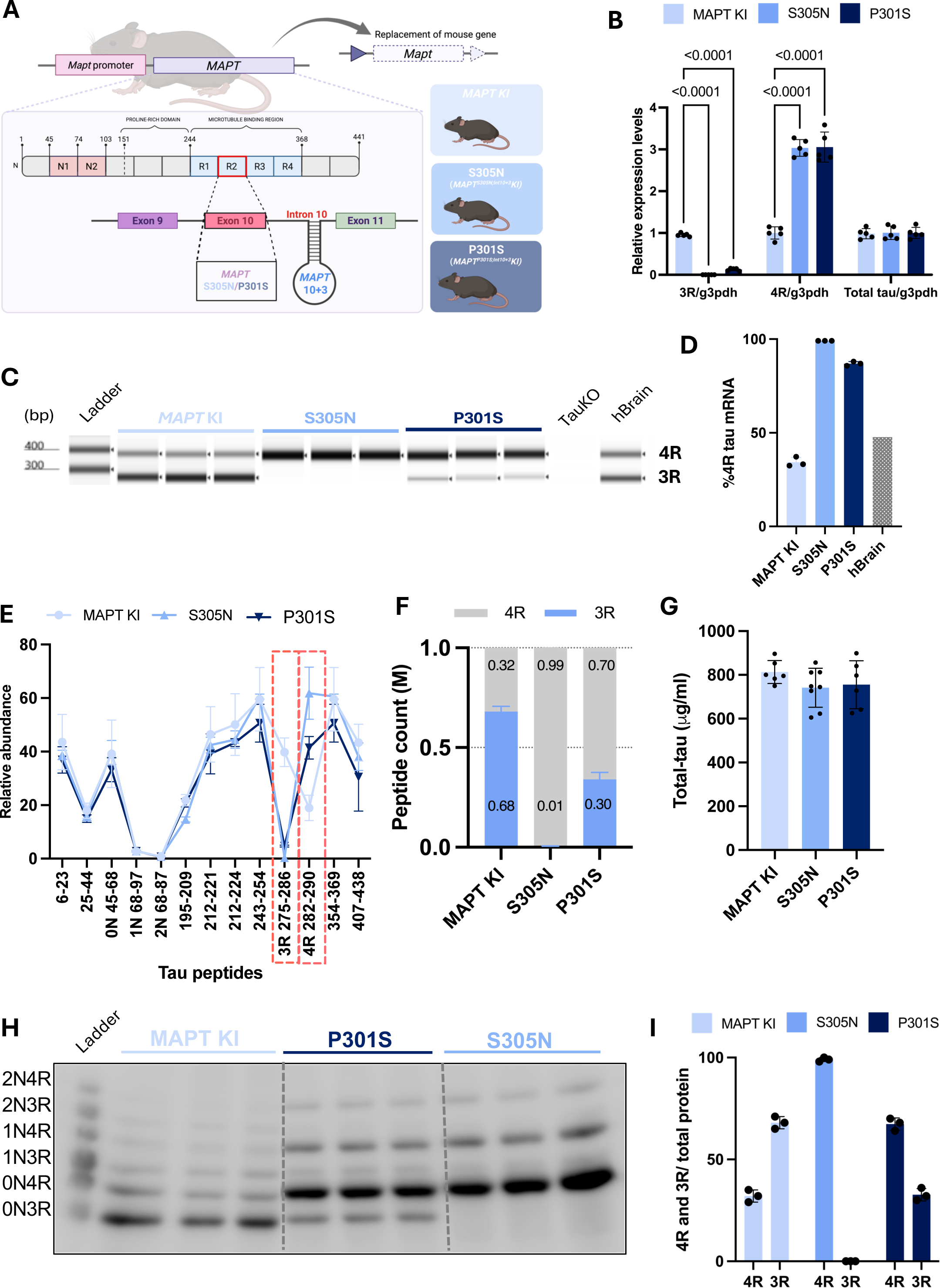
3R and 4R tau isoform ratio in S305N and P301S mice. (**A**) Schematic representation of generation of *MAPT* knock-in mouse models, showing the replacement of murine *Mapt* with human *MAPT* and insertion of S305N and intronic 10+3 or P301S and intronic 10+3. Schematic created using BioRender.com. (**B**) Real-time PCR results of 3R-tau, 4R-tau and total tau levels in *MAPT* KI, S305N and P301S KI mice (15M N=5 for each group; N=3 female and N=2 male) using specific primers validated in *Mapt* KO mice. Data represents mean±S.D. (**C**) TapeStation electrophoresis detecting PCR amplicons from 3R and 4R *MAPT* transcripts in *MAPT* KI mice, compared with mouse tau knockout (TauKO) and human brain samples. Bands corresponding to 3R and 4R tau are indicated by the ladder on the left. (**D**). Quantification of the % of 4R tau across genotypes. Samples from 15M KI mice (N=3 per group; N=2 females and N=1 male). Data represents mean±S.D. (**E**) Peptides profile along the tau chain from *MAPT* KI, S305N and P301S KI mice by LC–MS/MS analysis shown for individual tau peptides and 3R and 4R tau levels. (**F**) Results are expressed in L/H ratio. Red outline in E highlights 3R- and 4R-tau-specific tau peptides regions, respectively (N=5–9 for each group; *MAPT* KI: N=6 sex matched; S305N: N=9 of which N=4 female and N=5 male; P301S: N=5 of which N=3 female and N=2 male). (**G**) Total-tau ELISA level targeting the mid-region of the protein (N=6-8 each; MAPT KI N=4 female and N=2 males; S305N N=4 females and N=4 males; P301S N=3 females and N=3 males). Data represents mean±S.D. (**H**) Immunoblotting of tau detected by Tau13 antibody in *MAPT* KI, S305N and P301S KI mice (N=3 for each group, N=1 female, N=2 male) after alkaline phosphatase treatment and (**I**) quantification normalized to total protein. Data represents mean±S.D.

## Results

### *MAPT* mutation insertion in KI mice and splicing effects

To investigate the molecular basis of divergent tau disease phenotypes, we generated *MAPT* KI mouse lines expressing either *MAPT* S305N with the intronic 10+3 mutation (hereafter S305N) or *MAPT* P301S with the intronic 10+3 mutation (hereafter P301S). Both lines were established on a murine tau null background, with *MAPT* expression driven by the endogenous mouse promoter, to preserve physiologically relevant temporal and spatial expression (**Fig. 1A**). Real-time PCR (RT-PCR) revealed negligible expression of 3R tau RNA in S305N mice and low levels in P301S relative to the *MAPT* KI controls. Conversely, 4R-tau RNA was most abundant in S305N, followed by P301S. Total tau remained consistent across all genotypes (**Fig. 1B**). PCR analysis was consistent with these findings, showing ∼100% 4R in S305N, ∼86% 4R in the P301S and ∼34% in the *MAPT* KI line, confirming previous findings^26^ (**Fig. 1C-D**). At the protein level, mass-spectrometry (**Fig. 1E-F**) and western blotting (**Fig. 1H-I**) confirmed near-complete 4R isoform expression in S305N (∼99%) and a ∼70:30 ratio of 4R:3R in P301S. Total tau measured by ELISA showed no significant difference in tau levels across genotypes (**Fig. 1G**). These results demonstrate that mutations drive different splicing effects, with S305N expressing only 4R tau.

### S305N and P301S disrupt distinct cellular functions

Next, we asked whether *MAPT* mutations disrupt functional pathways in different ways. We first performed quantitative proteomic profiling of hippocampal proteins from *MAPT* S305N and P301S mice compared with *MAPT* KI controls. Gene set enrichment and network analysis of differentially abundant proteins revealed both shared and mutation-specific pathway alterations (**Fig. 2A; Fig. S1A-B**). Pathways related to cellular organization and ion transport were disrupted in both S305N and P301S mice. In S305N mice, proteins involved in cytoskeletal transport, protein localization, synaptic transmission, and vesicle trafficking were more affected, indicating synaptic and transport-related vulnerability, whereas P301S mice showed a distinct enrichment of changes in proteins involved in nucleotide metabolism and chromatin rearrangement-related proteins. To validate the proteomic findings at the transcriptional level, we performed single-nucleus RNA sequencing (snRNA-seq) and differentially expressed genes were subjected to pathway-level analysis and enrichment mapping. In a population of neurons that were especially vulnerable to tau pathology (hippocampal CA3 neurons), pathways related to synaptic transmission and microtubule polymerization were implicated in S305N mice; in P301S mice, mitochondrial metabolism and epigenetic regulators were implicated. These cross-platform concordances highlight mutation-specific perturbations in cellular processes and strengthen the link between the observed protein-level changes and transcriptional programs **(Fig. 2B; Fig. S2A-I**).

**Figure 2.**
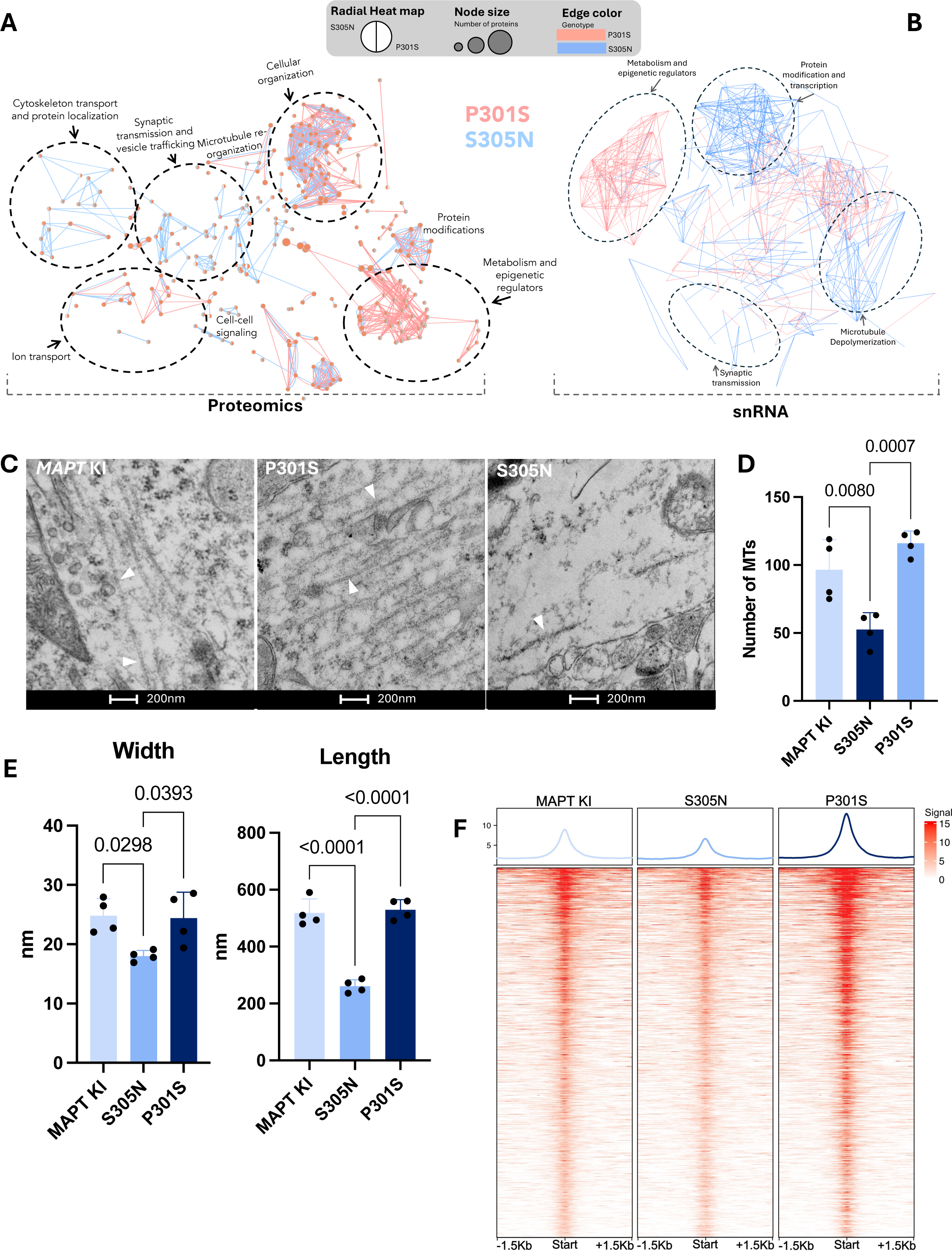
P301S and S305N affect different cellular pathways. (**A**) Differentially expressed proteins identified by TMT-based total proteomics dataset were analyzed for gene ontology enrichment using BiNGO in Cytoscape comparing S305N *vs MAPT* KI or P301S *vs MAPT* KI. Enrichment maps were generated with the EnrichmentMap app, displaying functional clusters of related GO terms. Blue edges indicate similarity between nodes in S305N and pink edges indicate similarity between nodes in P301S. A node cutoff of 0.89 and an edge cutoff of 0.75 were applied to define network connectivity. Dashed circles indicate manually annoted pathways based on GO terms (**B**) Significantly differentially expressed genes identified by snRNA sequencing were analyzed for gene ontology enrichment using BiNGO in Cytoscape comparing S305N *vs MAPT* KI or P301S *vs MAPT* KI. Enrichment maps were generated with the EnrichmentMap app, displaying functional clusters of related GO terms. Blue edges indicate similarity between nodes in S305N and pink edges indicate similarity between nodes in P301S. A node cutoff of 0.08 and an edge cutoff of 0.89 were applied to define network connectivity. Dashed circles indicate manually annoted pathways based on GO terms. (**C**) Representative electron microscopy (EM) images acquired from the pyramidal cell layer of the CA3 of the hippocampus from all genotypes. Ten images were collected per biological replicate (N=4 per genotype, all male). Image acquisition and analysis were performed blind to genotype and condition. Microtubules were quantified for number, diameter, and length within neuronal processes. White arrows point at microtubule. Quantification in (**D**) for number of microtubules along width and length (**E**) of microtubules in each genotype. Data represents mean±S.D. (**F**) Bulk ATAC-seq signal at the transcription start site (TSS - start, +/- 1.5Kb) across biological replicates for *MAPT* KI (N=4), S305N (N=3), and P301S (N=3) mice. Each row represents an average across replicates of a single site, and the mean peak intensity for each genotype is displayed above the corresponding heatmap.

To validate data implicating cytoskeletal disruption in S305N mice, we examined the ultrastructure of microtubules in CA3 pyramidal neurons by EM. Quantitative analysis revealed a significant reduction in microtubule number in S305N mice compared to both P301S and the *MAPT* KI. In addition, microtubules in S305N neurons were thinner and shorter, consistent with impaired microtubule stability or polymerization. P301S mice showed no significant alteration in microtubule morphology or abundance relative to *MAPT* KI (**Fig. 2C-E**).

Both proteomics and snRNA-seq revealed enrichment in P301S mice of pathways linked to metabolism, mitochondrial function, and energy production, as well as epigenetic regulation and RNA splicing (collectively referred to as metabolism and epigenetic regulators). These pathways are intrinsically interconnected: metabolic enzymes supply essential cofactors and substrates for chromatin modifications, while chromatin state influences splice-site recognition and alternative splicing. Thus, alterations in epigenetic regulators can directly impact RNA-binding and splicing proteins. To probe this relationship, we examined the status of the specific proteins involved. Data revealed downregulation of both metabolic and epigenetic regulators in P301S mice compared to S305N and control (**Fig. S1C**). We assessed chromatin accessibility in the mutant KI mice using bulk ATAC-sequencing and found a euchromatic profile in P301S, consistent with reduced activity of chromatin-modifying enzymes (**Fig. 2F**). A contributing factor could be the observed metabolic and mitochondrial alterations impacting the downstream SAM (S-adenosylmethionine) pathway, with reduced SAM potentially impairing methylation-dependent processes, including epigenetic and transcriptional regulation as well as RNA modifications. Consistent with this, SAM-related protein levels were downregulated in P301S mice. (**Fig. S1D**).

In summary, these data reveal divergent proteomic and pathway-level signatures that provide a mechanistic basis for *MAPT* mutation-driven neurodegeneration and functional deficits.

### Tau in S305N is hyperphosphorylated, but hypophosphorylated in P301S

Phosphorylation is among the most extensively studied post-translational modifications (PTMs), as tau isolated from *post mortem* AD or FTD brain tissue, is heavily hyperphosphorylated. However, the relationship between PTMs and tau aggregation – and the respective contributions of each to tau function and toxicity – remains incompletely understood and widely debated. To assess phosphorylation status, unbiased total phospho-proteomic analysis from hippocampi and posterior cortices of 15-month-old (M) *MAPT* KI, S305N and P301S was performed. ΔP analysis revealed that while the overall proportion of hyper- and hypo-phosphorylated proteins was similar across mutants, tau ranked as the second most hyperphosphorylated protein in S305N and the most dephosphorylated protein in P301S, relative to *MAPT* KI (**Fig. 3A**). To capture broad phosphorylation changes of tau, we built an eigenprotein score which confirmed a significant separation between groups, with S305N and P301S exhibiting distinct shifts compared to *MAPT* KI and to each other (**Fig. 3B**).

**Figure 3.**
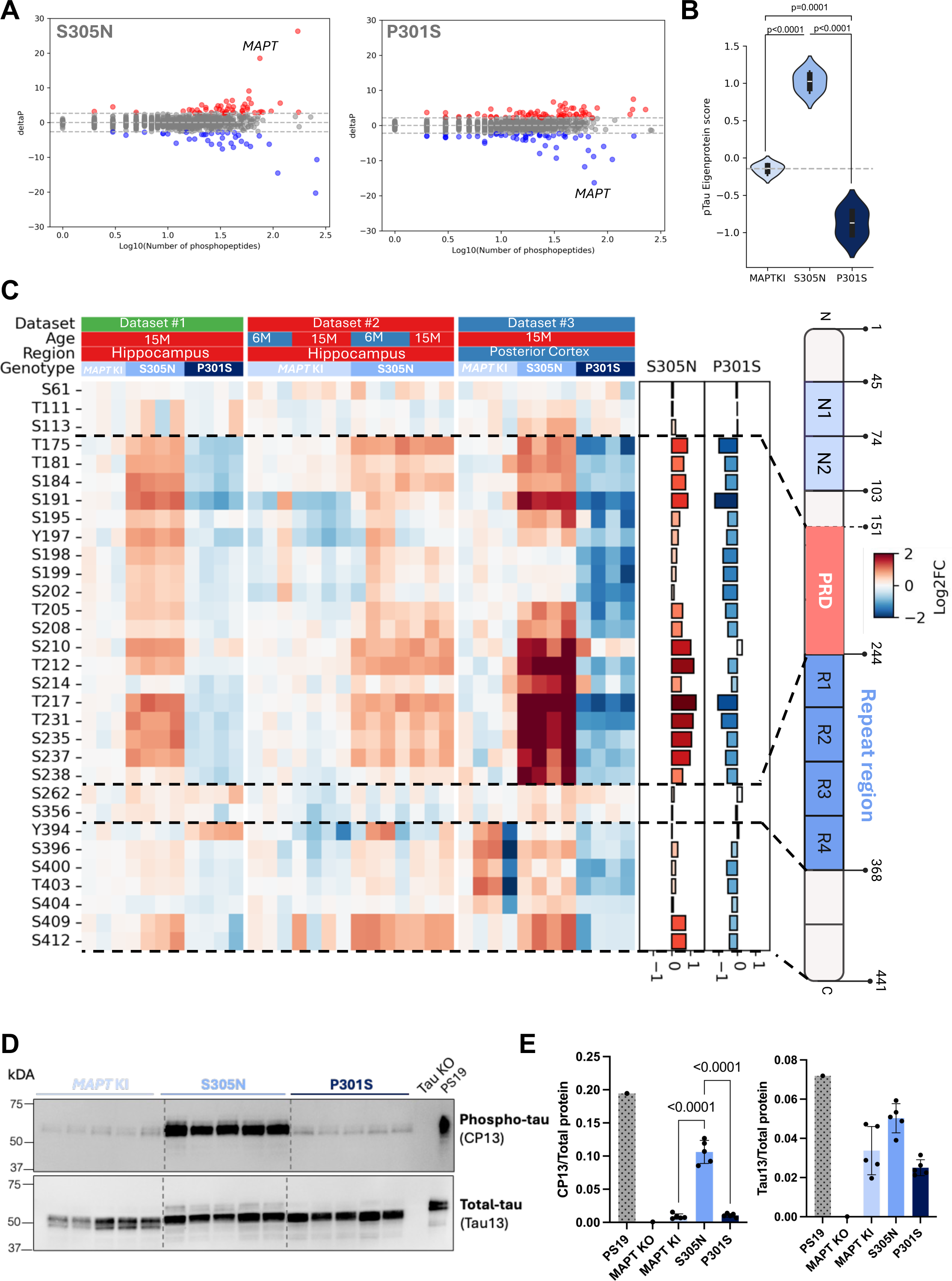
Hyperphosphorylation of tau in S305N and hypophosphorylation in P301S. (**A**) Phosphorylation status was calculated from TMT-based phospho-proteomics dataset 1. Overall phosphorylation status (ΔPs) of phosphoproteins was calculated as the sum of log₂(fold change;FC) values of all significantly changing phosphopeptides (p<0.05) compared with *MAPT* KI controls. Proteins with no phosphopeptides meeting the significance threshold were assigned ΔPs = 0. A stringent cut-off at ±3 standard deviations (σ) from the mean was applied to define cumulative phosphorylation changes, with ΔPs > 3σ representing hyperphosphorylation (red dots) and ΔPs < −3σ representing hypophosphorylation (blue dots). Samples included N=4 *MAPT* KI, S305N and P301S hippocampi at 15M. Phospho-tau data was normalised to total tau-peptides. (**B**) Eigenprotein values for phospho-tau–associated peptides were derived from principal component analysis of the TMT-based phospho-proteomics dataset 1. Scores are shown for *MAPT* KI controls, S305N, and P301S mice at 15 months. Group differences reflect the relative burden of tau phosphorylation across genotypes. Data are presented as median and IQR (dashed grey line crosses the median of *MAPT* KI control). (**C**) Heatmap representation of phosphorylated tau sites abundance across three independent TMT-based datasets: (#1) hippocampi from 15M *MAPT* KI, S305N, and P301S mice; (#2) hippocampi from 6M and 15M *MAPT* KI and S305N mice; and (#3) posterior cortical samples from 15M *MAPT* KI, S305N, and P301S mice. Values are expressed as log₂ fold change (log₂FC) relative to the *MAPT* KI controls from each dataset, with the colour scale ranging from red (increased phosphorylation) to blue (decreased phosphorylation). The accompanying barplots on the right depict the average of the fold change across all three datasets. (**D**) Immunoblotting of phosphorylated and total tau detected by CP13 and Tau13 antibodies in the RIPA fraction of hippocampi brain lysates from *MAPT* KI, S305N and P301S KI mice at 15M of age (N=5 for each group, N=3 female and N=2 males), along with positive control PS19 mice at 9M and Tau knock-out (KO) at 9M, and (**E**) quantification to total protein. Data represents mean±S.D.

To examine site-specific differences, we generated heatmaps of phospho-tau (p-tau) peptides normalized for *MAPT* KI, across three independent datasets **(Fig. 3C).** Across all datasets, the S305N mutant consistently exhibited significantly elevated phospho-tau levels compared to *MAPT* KI and P301S, particularly in the proline-rich domain (PRD; T151-S243 residues) and C-terminus region, possibly driven by S409 and S412. In contrast, the N-terminus and microtubule-binding domain (MTBR; 244-368) showed no statistical differences in phosphorylation levels across mutant lines and *MAPT* KI (**Fig. 3C and Fig. S3A**). This pattern was reproducible across both hippocampal and cortical datasets, indicating a robust and mutation-dependent phospho-tau signature. By contrast, the P301S mutant at 15M showed reduced or similar levels of phosphorylation at nearly all detected phospho-sites relative to *MAPT* KI. No significant enrichment was observed in any particular tau domain, and overall phosphorylation appeared less in P301S compared to both *MAPT* KI and S305N (**Fig. 3C and Fig. S3A**), in concordance with the ΔP analysis.

Targeted mass spectrometry on pre-selected tau peptides was performed to validate the phosphorylation status of specific tau epitopes across *MAPT* KI lines. This analysis revealed statistically increased levels of pT181, pT217, pT212/pT217, pT231, pT231+pS235, and pS396+pS404 in the S305N mice compared to *MAPT* KI (**Fig. S3B**).

Finally, western blot analysis was performed on hippocampal lysates from *MAPT* KI, S305N, and P301S mouse lines to assess total tau and p-tau levels. Total tau (Tau13 antibody) revealed comparable levels across all three genotypes. For p-tau, CP13 immunoblotting showed a significant increase in S305N compared to both *MAPT* KI and P301S (**Fig. 3D–E**). This pattern is consistent with our TMT-mass spectrometry data, which also demonstrated elevated S202 phosphorylation in S305N, while showing comparable levels between P301S and *MAPT* KI.

Together, these results demonstrate that the two mutations result in divergent tau phosphorylation signatures: S305N drives pronounced hyperphosphorylation, particularly in the PRD, while P301S at the same age drives global hypophosphorylation. To note, we define hyperphosphorylation and hypophosphorylation not to describe changes at individual peptide sites, but to reflect the cumulative phosphorylation state of the tau protein as a whole.

### Distinct Temporal and Conformational Profiles of Tau Pathology in S305N and P301S

We next compared the distribution and progression of tau pathology in S305N and P301S mice. Immunohistochemistry was performed on hippocampal sections from *MAPT* KI, S305N, and P301S mice at 15 and 30M, using antibodies against phosphorylated tau (AT8/CP13), oligomeric tau (TOC1), and misfolded/aggregated tau (MC1). At 15M, *MAPT* KI mice showed no detectable tau staining. S305N mice were positive for AT8 and CP13, showing perinuclear and diffuse accumulation of phosphorylated tau, but were negative for TOC1 and MC1, indicating the absence of oligomeric or conformationally altered tau species. P301S mice showed negligible staining with any antibody at this age. By 30M, P301S mice had developed focal pathology marked by AT8, CP13, TOC1, and MC1 positivity, reflecting the emergence of both phosphorylated and conformationally altered tau. Staining was restricted to sparse individual neurons, unlike the diffuse labeling seen in S305N mice, which at 30M continued to show only diffuse AT8 and CP13 immunoreactivity. *MAPT* KI mice remained essentially negative at all ages (**Fig. 4A, Fig. S3C**)

**Figure 4.**
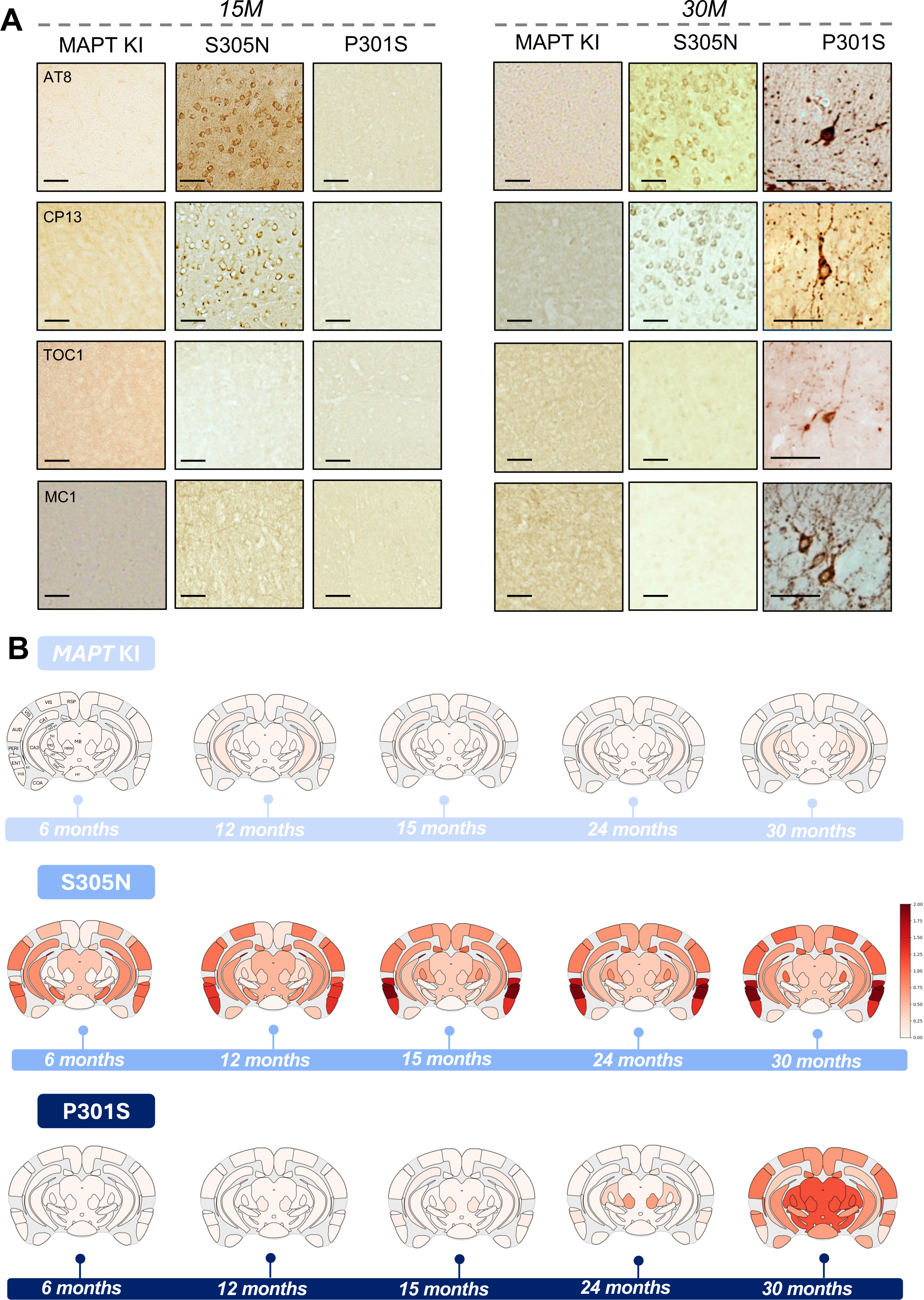
Spatial and temporal profiling of tau pathology in KI mutants. (**A**) Tau immunostaining detected by AT8 (phosphorylated tau), CP13 (phosphorylated tau), TOC1 (oligomeric tau) and MC1 (conformationally abnormal tau) antibodies in the brains of *MAPT* KI, S305N and P301S at 15M (left panel) and 30M (right panel). Scale bar=50µm. (**B**) Heatmaps showing AT8 signal intensity from coronal sections of *MAPT* KI, S305N and P301S at 6, 12, 18, 24 and 30M. N=4/5 cases per age, except P301S at 30M (N=5/7). Bregma is located at -3.07mm (according to Paxinos and Franklin mouse brain atlas). Labels of heatmap found in 6M *MAPT* KI: RSP: Retrosplenial area, VIS: Visual area (visual cortex), AUD: Auditory area (auditory cortex), PERI: Perirhinal area, ENT: Entorhinal area, CA1: Field CA1 of hippocampus, CA3: Field CA3 of hippocampus, SUB: Subiculum, DG-po: Dentate gyrus, polymorph layer (hilus), TH: Thalamus, MB: Midbrain, ec: External capsule, MRN: Midbrain raphe nuclei, SNc: Substantia nigra, compact part, MG: Medial geniculate complex, SCm: Superior colliculus, motor-related, HY: Hypothalamus, COA: Cortical amygdalar area, PIR: Piriform area (piriform cortex).

To track spatiotemporal progression, we generated semi-quantitative heatmaps of AT8 staining across brain regions in animals aged 6, 12, 15, 24 and 30M (**Fig. 4B**). Coronal sections spanning the hippocampus, auditory, visual, ectorhinal and entorhinal cortices, thalamus and hypothalamus, amygdala and midbrain regions were scored for immunoreactivity to AT8. Heatmap representation of *MAPT* KI mice revealed no marked spatiotemporal differences in pathology, apart from a slight but variable increase in phosphorylated tau signal in the polymorphic layer of the hippocampus from 6 to 12M, after which staining intensity remained consistent across ages and brain regions. In S305N mice, tau pathology was first detectable at 6M, with a modest increase at 15M, after which the distribution remained consistent through 30M. The pathology was predominantly localized to the hippocampus, entorhinal cortex, and cortical regions, with low signal in the subcortical regions, and was characterized by diffuse, widespread labeling and perinuclear phosphorylated tau signal. In contrast, P301S mice exhibited no detectable pathology until 24M, at which point sparse, focal neuronal staining emerged, affecting individual neurons in the midbrain, with no visible diffuse signal. By 30M, sparse neuronal tau staining occurred in cortical areas and hippocampus, but was strongest in thalamus and midbrain. The extent and distribution of pathology in P301S mice at 30M were variable across individuals (N=7) (**Fig. 4B; Fig. S4**).

These findings highlight divergent trajectories of tau pathology: S305N mice develop early, diffuse accumulation of hyperphosphorylated, conformationally normal tau, whereas P301S mice show late-onset, focal pathology associated with late-stage accumulation of phospho-epitopes and a change in conformation and assembly. These differences in timing, distribution, and molecular profile point to distinct mechanisms of tau accumulation and spread.

### Tau in P301S is seed-competent and fibrillar

To assess whether tau from P301S or S305N KI mice undergoes a conformational change enabling prion-like seeding^27^, we employed tau biosensor cell lines expressing either the repeat domain of tau with the P301S mutation fused to CFP and YFP (P301S biosensor) or an in-house version with the S305N mutation fused to YFP (S305N biosensor)^26^. In the P301S biosensor line, TBS-soluble extracts from the posterior cortex of 15M P301S mice (a region that showed no histological pathology at the same age) induced seeding activity. In contrast, TBS extracts from age-matched S305N KI mice and *MAPT*KI controls failed to elicit detectable seeding, indicating a lack of conformationally abnormal tau in these genotypes at this age (**Fig. 5A**). The same result was observed in the S305N biosensor, with P301S but not the S305N and *MAPT* KI lines showing the presence of tau seeds (**Fig. S5A**).

**Figure 5.**
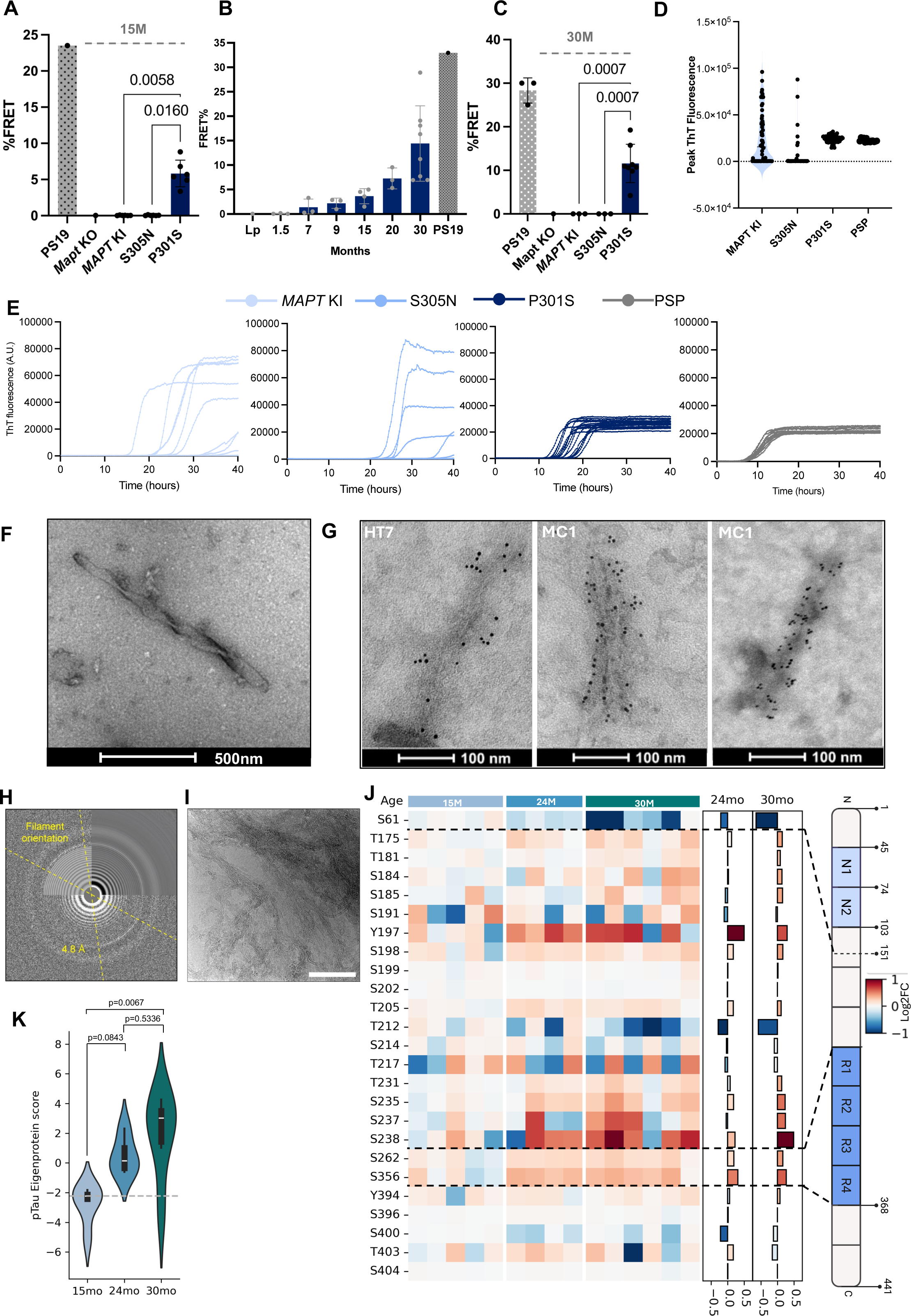
Tau in P301S is seed-competent, fibrillar and becomes phosphorylated. (**A**) Percentage FRET of tau seeding activity on P301S biosensor in brain lysates from *MAPT* KI, S305N and P301S mice at 15M (N= 6 for each group, N=3 females and N=3 males) and PS19 mouse at 9M. Data represents mean±S.D. (**B**) Percentage FRET of tau seeding activity on P301S biosensor in brain lysates from P301S mice at different months: 1.5 (N=3), 7 (N=3), 9 (N=3), 15 (N=4), 20 (N=3), 30 (N=8), compared to PS19 mouse at 9M and lipofectamine negative control (“Lp”). Data represents mean±S.D; (**C**) Percentage FRET of tau seeding activity on P301S biosensor in brain lysates from *MAPT* KI, S305N and P301S mice at 30M (N=8 for each group, N=4 females and N=4 males) compared to PS19 mouse at 9M and MaptKO mouse at 9M; (**D**) RT-QuIC aggregates at peak ThT fluorescence (within 40h time) from the combination of 16 replicates per biological sample from 15M posterior cortex from *MAPT* KI (N=4; sex matched), S305N (N=3; N=2 female and N=1 male), P301S (N=4; sex matched) and PSP human brain as positive control (N=1). (**E**) Raw RT-QuIC ThT reactions from posterior cortex of mice at 15 months, *MAPT* KI (N=4), S305N (N=3) and P301S (N=4) and positive control PSP human brain homogenates (N= 1). N=1 is represented in this figure (the additional Ns can be found in Supplemental Fig. 2). (**F**) Representative negative-stain EM images of sarkosyl-insoluble tau fibrils isolated from whole-brain homogenates of 30M P301S mice (N=5). (**G**) Immunogold labelling (tau antibodies MC1 and HT7 on 6nm gold beads) of sarkosyl-insoluble extracted fibrils from whole-brain of 30M P301S mice. (**H**) Fast Fourier transform (FFT) analysis on cryoEM images showing characteristic 4.8 Å meridional reflection and (**I**) raw cryoEM images from 30M P301S sarkosyl-insoluble extraction showing clusters of filaments. Scale bar represents 100nm. Raw images showing untwisted and thicker filaments are found in supplemental Fig. 2E-F. (**J**) Heatmap representation of phosphorylated tau sites from DIA-acquired phospho-proteomics dataset comparing hippocampi from 15M, 24M and 30M P301S mice. Values are expressed as log₂ fold change (log₂FC), with the colour scale ranging from red (increased phosphorylation) to blue (decreased phosphorylation). The barplot on the right depicts the average of the fold change at 24M and 30M; (**K**) Eigenprotein values for phospho-tau–associated peptides were derived from principal component analysis from DIA-acquired phospho-proteomics dataset. Scores are shown for P301S mice at 15M, 24M and 30M. Group differences reflect the relative burden of tau phosphorylation across ages. Data are presented as median and IQR (dashed grey line crosses the median of P301S at 15M). Statistical comparisons were performed using one-way ANOVA with Tukey’s Honestly Significant Difference.

To define the temporal onset of seeding activity in P301S mice, we conducted a longitudinal analysis at 1.5, 7, 9, 15, 20 and 30M. Seeding activity was first detectable at approximately 9M, which progressively increased thereafter, suggesting that tau conformers accumulate over time in this model. No, or inconsistent seeding was observed at earlier timepoints (1.5-7 months) (**Fig. 5B**). No seed-competent tau was detected at 30M in *MAPT* KI and S305N lines (**Fig. 5C**), including in the S305N biosensor line (**Fig. S3B**).

To further evaluate whether tau in P301S KI mice is seed-competent, we employed a 4R-tau-specific real-time quaking-induced conversion (RT-QuIC) assay, which quantifies the ability of tau to nucleate and polymerize monomeric tau into fibrils, as measured by thioflavin T (ThT) fluorescence^28^. PBS homogenates from the posterior cortex of 15M P301S mice consistently induced fibrillization, with ThT fluorescence amplitudes clustering within a defined range, and showing comparable peak ThT fluorescence and half-times (t_1/2_) across technical replicates, indicating the presence of conformationally stable tau seeds (**Fig. 5D-E; Fig. S5C-D**). This result was similar for the positive control (human progressive supranuclear palsy brain extract). In contrast, homogenates from *MAPT* KI controls and S305N mice exhibited delayed and variable fluorescence signals, characterized by inconsistent amplitudes, peak ThT fluorescence and t_1/2_ values (**Fig. 5D-E; Fig. S5C-D**). These features are indicative of stochastic, spontaneous nucleation events rather than templated polymerization. These data confirm that, unlike *MAPT* KI and S305N mice, P301S mice harbor tau conformers with reproducible and templating-capable conformations.

To investigate whether aging promoted the formation of higher-order tau assemblies in *MAPT* KI P301S mice, we examined the sarkosyl-insoluble fraction from whole brains of 30M animals by immunogold labelling and negative-stain transmission electron microscopy (EM). Ultrastructure analysis revealed fibrillar material in P301S mice but not in *MAPT* KI and S305N (**Fig. 5F; Fig. S5E**). Immunogold labelling on P301S confirmed that these filaments were positive for HT7 (recognizing mid-region tau) and MC1 (**Fig. 5G**). We next imaged P301S 30M sarkosyl-insoluble tau preparations by cryoEM which revealed untwisted filaments ranging from 7 to 16 nm in width, and thicker (13-38 nm wide) filaments with apparent 280-350 nm crossover distance (**Fig. S5F-G**). 2D class averages from cryoEM data revealed insufficient twist to solve the structure but fast Fourier transform (FFT) analysis of some filaments revealed a characteristic 4.8 Å meridional reflection, indicative of cross-β sheet spacing consistent with tau filament architecture (**Fig. 5H-I**). This is, to our knowledge, the first report of EM-validated tau filaments in a non-overexpressing mouse model of tauopathy.

To determine whether the phosphorylation profile of tau evolves with the transition to a fibrillar state, we performed phospho-proteomic analysis on TBS-soluble brain extracts from P301S KI mice at 15, 24, and 30M. Plotting PC1 eigenprotein scores for each timepoint demonstrated an age-dependent trajectory, with 24M samples occupying an intermediate state between 15M and 30M; only the 15M *versus* 30M comparison reached statistical significance, consistent with progressive tau phosphorylation with age (**Fig. 5K**). A trend for increased phosphorylation was noted in the PRD and MTBR at 24M, which became more pronounced at 30M. Whilst the C-terminus of tau did not show differences in phosphorylation levels, the N-terminus was significantly decreased in the 30M time-point (**Fig. 5J and Fig. S5H**). These changes in phosphorylation coincided temporally with the emergence of tau filaments, suggesting that altered post-translational modification may either facilitate or result from tau aggregation in old P301S mice.

Overall, these findings show that tau in *MAPT* KI P301S mice is hypophosphorylated compared to *MAPT* KI and S305N, yet this state is compatible with the formation of seed-competent tau. With age, *bona fide* tau filaments are observed. Fibrillization is accompanied by a shift from hypophosphorylation toward progressive phosphorylation of tau, linking phosphorylation state with aggregation status and reflecting the hyperphosphorylated, fibrillar tau seen in human tauopathies at end-stage. In contrast, tau in S305N mice remained soluble with no detectable seeds or fibrils, even in aged mice, again underscoring a distinct pathogenic trajectory in the two mutants.

### β-Amyloid accelerates neuropathology

To explore how β-amyloid interacts with tau in S305N *versus* P301S mice, we crossed the *MAPT* KI mutant lines with the APP^NL-G-F^ KI line (NLGF) which develops β-amyloid plaques from 2M^29^. This generated NLGF-*MAPT* KI, NLGF-S305N, and NLGF-P301S lines.

To assess whether the accumulation of β-amyloid accelerated tau neuropathology, we performed immunohistochemistry for p-tau (CP13) on 15M mice. As expected, p-tau accumulating in dystrophic neurites around plaques in NLGF-positive mice was seen in all mice. NLGF-*MAPT* KI mice showed negligible somatic tau pathology at this age. NLGF-S305N mice displayed a pattern of somatic CP13 immunoreactivity comparable in distribution and intensity to that observed in S305N mice: TOC1 and MC1 were negative (**Fig. 6A-B; Fig. S6A**). In contrast, NLGF-P301S mice exhibited marked enhancement of somatic tau pathology compared to P301S at the same age (**Fig. 6C**). Immunopositive neurons were sparse but intensely labelled with CP13, AT8 (phosphorylated tau found in pretangles and mature tangles), MC1, PHF1 (phosphorylated tau indicative of more mature tangles), and Gallyas silver stain (indicative of insoluble, fibrillar inclusions) (**Fig. 6C**).

**Figure 6.**
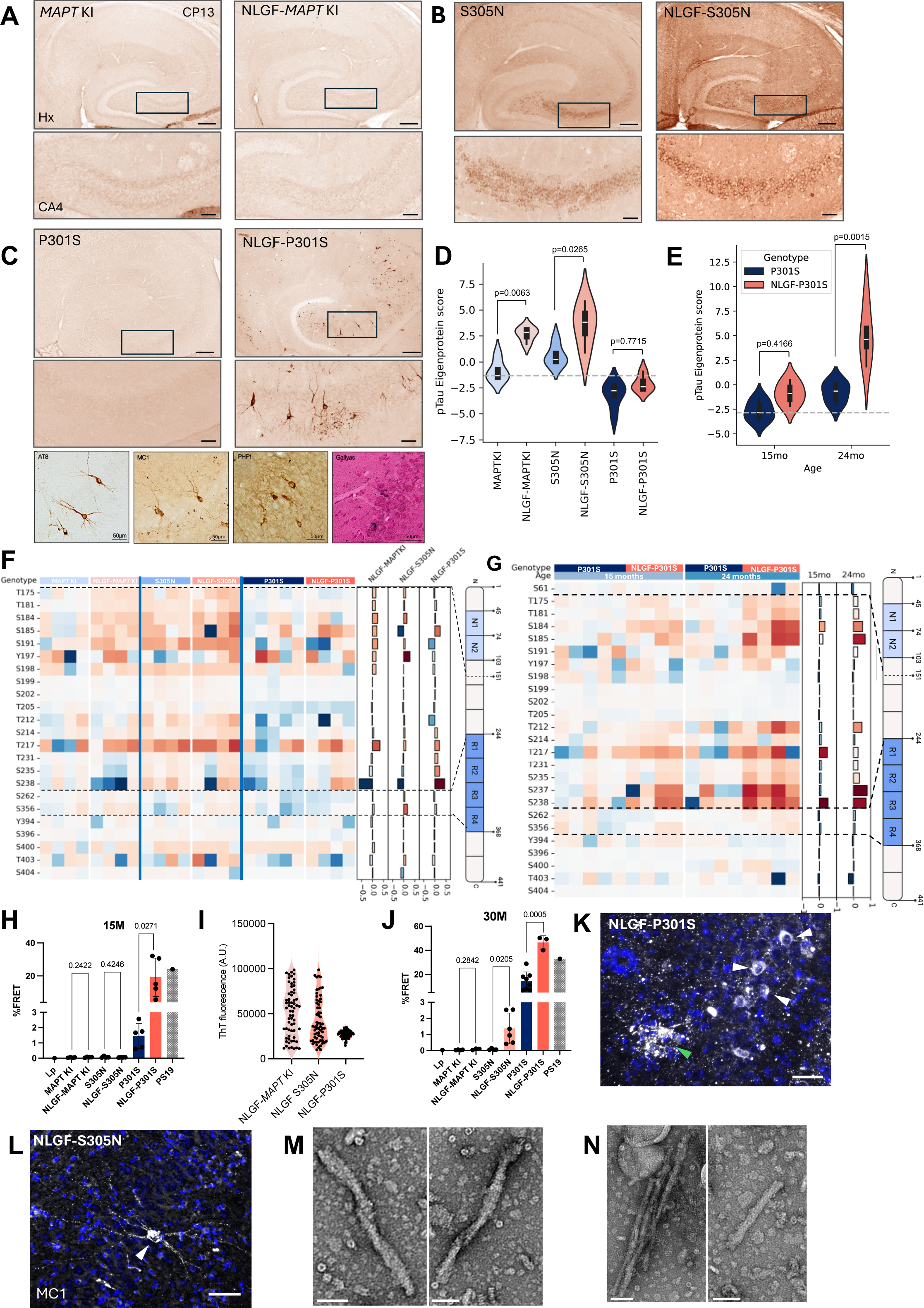
β-amyloid exacerbates seed-competent tau formation. Representative overview images of immunostaining of phosphorylated tau detected by CP13 antibody comparing the hippocampus and CA4 between 15M *MAPT* KI and NLGF-*MAPT* KI (**A**), 15M S305N and NLGF-S305N (**B**) and 15M P301S and NLGF-P301S (**C**). Scale bars represent 200 µm (hippocampus, *i.e.* upper panel) and 100 µm (CA4, *i.e.* lower panel), unless specified (lower focused panel from NLGF-P301S neurons = 50µm). Black box in the hippocampus image indicates the zoomed in area of the CA4 (below) (**D**) Eigenprotein values for phospho-tau–associated peptides were derived from principal component analysis of the DIA-acquired phospho-proteomic dataset. Scores are shown comparing *MAPT* KI *vs* NLGF-*MAPT* KI controls (N=4; N=2 male and N=2 females), S305N *vs* NLGF-S305N (N=4; N=2 male and N=2 females), and P301S (N=5; N=2 male and N=3 females) *vs* NLGF-P301S (N=4; N=2 male and N=2 females) mice at 15M. Group differences reflect the relative burden of tau phosphorylation across genotypes. Data are presented as median and IQR (dashed grey line crosses the median of *MAPT* KI control). Statistical comparisons were performed using one-way ANOVA with Tukey’s Honestly Significant Difference. (**E**) Eigenprotein values for phospho-tau–associated peptides were derived from principal component analysis of DIA-based phospho-proteomics dataset. Scores are shown comparing 15M P301S (N=5; N=2 males and N=3 females) *vs* NLGF-P301S (N=4; N=2 male and N=2 females) and 24M P301S *vs* NLGF-P301S (N=4; N=2 male and N=2 females). Group differences reflect the relative burden of tau phosphorylation across genotypes. Data are presented as median and IQR (dashed grey line crosses the median of P301S at 15M). Statistical comparisons were performed using one-way ANOVA with Tukey’s Honestly Significant Difference. (**F**) Heatmap representation of phosphorylated tau sites from DIA-based phospho-proteomics dataset comparing hippocampi from *MAPT* KI *vs* NLGF-*MAPT* KI controls (N=4; N=2 male and N=2 females), S305N *vs* NLGF-S305N (N=4; N=2 male and N=2 females), and P301S (N=5; N=2 male and N=3 females) *vs* NLGF-P301S (N=4; N=2 male and N=2 females) mice at 15M. Values are expressed as log₂ fold change (log₂FC), with the colour scale ranging from red (increased phosphorylation) to blue (decreased phosphorylation). The barplot on the right depicts the average of the fold change of NLGF-*MAPT* KI (compared to *MAPT* KI), NLGF-S305N (compared to S305N) and NLGF-P301S (compared to P301S). (**G**) Heatmap representation of phosphorylated tau sites from DIA-based phospho-proteomics dataset comparing hippocampi from P301S and NLGF-P301S at 15M (N=5; N=2 males and N=3 females) and 24M (N=4; N=2 male and N=2 females). Values are expressed as log₂ fold change (log₂FC), with the colour scale ranging from red (increased phosphorylation) to blue (decreased phosphorylation). The barplot on the right depicts the average of the fold change of NLGF-P301S (compared to the respective age-related P301S). (**H**) Percentage FRET of tau seeding activity measured in P301S biosensor cells exposed to brain lysates from 15M *MAPT* KI, S305N, P301S, and their corresponding 15M NLGF lines. Each group included N=4 mice (2 females, 2 males), except P301S and NLGF-P301S, which included N=5 mice (3 males, 2 females) and was compared to PS19 positive control at 9M and lipofectamine only negative control. Data represent mean±S.D and statistics performed as Welch’s t-test comparing KI to the corresponding NLGF-KI. Lp=lipofectamine only. (**I**) RT-QuIC aggregates ThT maxima from the combination of 16 replicates per biological sample from 15M posterior cortex from NLGF-*MAPT* KI (N=4), NLGF-S305N (N=4) and NLGF-P301S (N=4). (**J**) Percentage FRET of tau seeding activity measured in P301S biosensor cells exposed to brain lysates from 30M sex-matched *MAPT* KI (N=4), S305N (N=4), P301S (N=8), and 30M NLGF-*MAPT* KI (N=3, 2 females and 1 male), NLGF-S305N (N=6) and NLGF-P301S (N=3, 2 females and 1 male). FRET-signal was compared to PS19 positive control at 9M and lipofectamine only negative control. Data represent mean±S.D and statistics performed as Welch’s t-test comparing KI to the corresponding NLGF-KI. Lp=lipofectamine only. (**K**) Representative overview images of immunostaining of conformational-specific tau antibody (MC1) in the hippocampus from 30M NLGF-P301S. Scale bar represents 50µm. White arrows point at neurons, green arrow points at dystrophic neurite. (**L**) Representative overview images of immunostaining of conformational-specific tau antibody (MC1) in the hippocampus from 30M NLGF-S305N. Scale bar represents 50µm. White arrows point at neuron. (**M**) Representative negative-stain EM images of sarkosyl-insoluble tau fibrils isolated from whole-brain homogenates of 24M NLGF-P301S mice (N=2). Scale bar represents 50nm. (**N**) Representative negative-stain EM images of sarkosyl-insoluble tau fibrils isolated from whole-brain homogenates of 30M NLGF-S305N mice (N=2). Scale bar represents 50nm.

To better quantify the effects of β-amyloid accumulation on phospho-tau accumulation, we performed an unbiased phospho-proteomic analysis of hippocampal extracts. Focusing on tau peptides, at 15M a significant increase was seen only in NLGF-*MAPT* KI and NLGF-S305N, relative to *MAPT* KI and S305N mice respectively. This modest increase in phospho-tau in NLGF-expressing mice likely reflects the local accumulation in dystrophic neurites around plaques rather than global biochemical shifts. Although a trend was observed for NLGF-P301S relative to P301S mice, this did not reach significance (**Fig. 6D**). However, by 24M, when P301S mice began to develop hyperphosphorylated tau accompanied by fibrillar inclusions, NLGF-P301S mice exhibited a significant shift in eigenprotein scores relative to age-matched P301S (**Fig. 6E**). Overall, these shifts occurred without detectable change in total tau abundance, as confirmed by western blot (**Fig. S6B-C**). In NLGF-*MAPT* KI and NLGF-S305N mice compared to the *MAPT* KI and S305N counterparts, increased phosphorylation was observed at sites in the PRD and in the MTBR for NLGF-S305N only (**Fig. 6F**). In NLGF-P301S, site-specific analysis revealed early and progressive changes. At 15 months, phosphorylation was elevated in the PRD, including T217, in NLGF-P301S mice compared to P301S. By 24 months, additional increases were observed at S184, T212, T237, and T238 (**Fig. 6G)**.

### β-Amyloid accelerates seed-competent tau formation

To assess the impact of β-amyloid on pathways resulting in tau seed formation, we performed the FRET-based biosensor assay using hippocampal lysates from 15M mice. No seeding activity was detected in lysates from NLGF-*MAPT* KI or NLGF-S305N mice (**Fig. 6H**) compared to respective *MAPT* KI or S305N mice. In contrast, NLGF-P301S mice displayed a significant increase in seed-competent tau relative to P301S (**Fig. 6H**). To confirm the results of the biosensor assay, we performed the RT-QuIC assay. Samples from NLGF-*MAPT* KI and NLGF-S305N mice exhibited delayed, low-amplitude, and highly variable ThT responses, with high inter-replicate variance in t_1/2_ values (**Fig. 6I and Fig. S6D-E**), indicative of spontaneous nucleation events, and therefore absence of structured, seed-competent tau assemblies. Homogenates from NLGF-P301S mice consistently induced robust ThT fluorescence, with fluorescence amplitudes clustering within a narrow dynamic range and half-times (t_1/2_) of aggregation tightly grouped (**Fig. 6I and Fig. S6D-E**), indicating the presence of conformationally-stable tau seeds capable of templated fibrillization.

To assess whether tau in non-seeding lines could evolve over longer timeframes, we extended our analysis to aged cohorts. At 30M NLGF-*MAPT* KI continued to show no seeding-activity. Both P301S and NLGF-P301S mice showed robust seeding activity, with NLGF-P301S exceeding the seeding levels of the PS19 positive control (**Fig. 6J**). NLGF-S305N mice, which were previously devoid of detectable seeding activity, also developed significant tau seeding activity at this age (**Fig. 6J**). This observation was consistent in the S305N biosensor line (**Fig. S6F**)

### β-Amyloid promotes tau fibril formation

Given the robust seeding activity observed in NLGF-P301S mice, we next investigated whether this translated into increased tau filament formation. At 30M, NLGF-P301S and NLGF-S305N showed positive MC1-neuronal staining (**Fig. 6K-L**). Immunogold EM of sarkosyl-insoluble extracts from 24M NLGF-P301S brains confirmed the presence of HT7 and MC1-positive fibrils (**Fig. S6G**). Subsequent negative stain EM revealed fibrils in NLGF-P301S brains at 24M (**Fig. 6M**) that were more readily detectable than the sparse fibrils observed in age-matched P301S mice. Of note, NLGF-S305N mice at 30M also showed negative stain EM tau filaments (**Fig. 6N**) demonstrating that tau filaments could form in these mice, but only with extreme aging and when exposed to elevated β-amyloid.

In summary, β-amyloid strongly potentiates fibrillar tau formation in P301S mice and, with prolonged aging, can drive fibrillization in S305N mice.

## Discussion

By leveraging physiologically relevant *MAPT* KI mice we demonstrate that distinct *MAPT* mutations drive fundamentally different trajectories of early tau pathogenesis: one (S305N) associated with tau hyperphosphorylation, disrupted cytoskeletal and synaptic dynamics; the other (P301S) associated with seed formation and disrupted nucleotide metabolism. These findings challenge the notion of a linear model of tau pathogenesis and argue for mechanistically divergent routes to neurodegeneration in the tauopathies.

Despite overt hyperphosphorylation in the proline rich domain (PRD), tau in 15M old S305N mice was soluble, devoid of seeds and fibrils. At the same age, there was surprisingly less phosphorylated tau in P301S mice relative to S305N mice and even to control *MAPT* KI, yet tau was robustly seed-competent. With extreme aging, tau in S305N mice did not change significantly but in the P301S line it became hyperphosphorylated (more in the MTBR than for S305N), and *bona fide* fibrils were observed that were comparable to the untwisted filaments described in P301L human mutation carriers^30^. The initial decoupling of hyperphosphorylation from aggregation contrasts with *in vitro* or cell-free systems that propose that phosphorylation is a prerequisite for fibrillization. Phosphorylation has been proposed to destabilize the native “paperclip” structure of tau, or mobilize tau from microtubules and promote aggregation^31–33^. Conversely, other phosphorylation events have been shown to suppress fibrillization^34,35^. The link between hyperphosphorylation and reduced tau-microtubule binding, particularly across the interaction site spanning aa202-365^36^ has been well-documented. The PRD segment (202–243), heavily phosphorylated in S305N mice, includes T231^37^ and S235^36^ sites, which are known to weaken tau’s ability to assemble and stabilize microtubules. In general, the *MAPT* KI mice will be valuable tools to work out the complex relationship between phosphorylation and aggregation that can only be fully reproduced in a physiologically relevant *in vivo* system. Finally, loss of normal tau function with microtubules is consistent with reports linking microtubule dysfunction to spine retraction and synapse loss^38–40^, which we previously showed as heavily affected in S305N^26^.

P301S mice demonstrated an enrichment in pathways related to metabolism, epigenetic regulators and RNA processing. Altered metabolism can disrupt epigenetic regulation by histone-and DNA-modifying enzymes, suggesting that the tau-driven euchromatin profile may arise downstream of metabolic changes^41^. Overall, we observed not only a reduction in the enzymes responsible for SAM biosynthesis but also a downregulation of multiple methylation “writers,” including RNA-specific methyltransferases, pointing toward a broader dysregulation of protein methylation, both at the transcriptomic and proteomic level, consistent with the euchromatic profile. Among the chromatin regulators, the reduction of NCOR2 was particularly striking given its established role in controlling chromatin accessibility around transcriptional start sites^42^. The enrichment of RNA splicing and processing pathways may reflect the P301S-specific euchromatic state, which accelerates RNA Pol II transcription and alters chromatin marks, thereby disrupting spliceosome recruitment at exon–intron junctions^43^. These findings converge with studies in drosophila, mouse, and Alzheimer’s disease human brain tissue, showing that tau aggregation induces heterochromatin loss and chromatin relaxation^44^, and with epigenome-wide analyses in aging human brains linking tau burden to altered histone acetylation, reflecting large-scale chromatin reorganization^45,46^. Thus, epigenomic remodeling emerges as a distinguishing feature of aggregation-driven tauopathy.

β-Amyloid showed a strong synergistic effect on tau pathogenesis. In NLGF-P301S mice, β-amyloid strongly enhanced both seeding and fibril formation pathways, leading to levels of insoluble tau that were higher than the P301S transgenic model (PS19). β-Amyloid did not alter the phosphorylation profile in NLGF-S305N mice significantly but in extreme old age, fibrillar tau was observed, presumably formed either by exacerbating fibrillization of some slow-developing sub-detection tau seeds, or by inducing *de novo* synthesis. Although β-amyloid accumulation has been shown previously to accelerate and exacerbate the accumulation of aggregated tau *in vivo*^25,47^, demonstrating that the effect is most likely due to enhancement of pathway/s leading to seed formation rather than tau hyperphosphorylation has not been possible previously. This distinction is critical for modelling the interaction, so that therapeutic targets aiming to decouple amyloid from tau can be identified and developed.

Humans with FTD show a high degree of clinical heterogeneity. Although very few studies have been performed in *ante mortem* carriers of *MAPT* mutations, a couple of studies suggest that divergence between splice-shifting and aggregation-promoting mutations occurs in humans. Young et al.^12^ demonstrated that *MAPT* 10+16 mutation carriers (similar in effect to S305N) exhibited a temporal lobe-predominant atrophy profile, whereas P301L mutation carriers (similar to P301S) show additional fronto-occipital involvement. This is in line with our work showing a divergent pathology distribution pattern. In another study^11^ donors expressing splice-shifting mutations (S305I, 10+16) were shown to have higher levels of 4R tau in their CSF compared to donors expressing exon 10 mutations associated with aggregation (*e.g.,* P301L). Of note, the splice-shifting human mutation carriers also showed relatively low levels of tau aggregates. Divergence is also seen at the structural level as cryo-EM studies show that fibrillar tau extracted from 10+3/16 mutations adopts a conformation that is markedly different from P301L/T carrier tau^48,49^. As human data is usually *post mortem*, and from a late- or end-stage of the disease, the mice will be useful additions to our toolbelt to understand how distinct tau profiles defined by isoform composition, phosphorylation state, conformation, and the selective disruption of specific cellular pathways, can contribute to clinical heterogeneity.

The observation of divergent mechanistic pathways has important implications for therapeutic development. Strategies aimed at preventing or reducing tau aggregate formation, may be effective for seed-driven tauopathies, but ineffective for tauopathies dominated by soluble, hyperphosphorylated tau toxicity, which may respond better to targeting cytoskeletal stability or reducing phosphorylation. Our data further argue for patient stratification approaches based on biochemical and conformational signatures of tau in peripheral biofluids, rather than treating tauopathies as a single disease entity.

Our study underscores the value of *MAPT* KI models for dissecting initiating events associated with tau pathogenesis, and for better defining the role of triggers such as β-amyloid. With their hyperphosphorylated, fibrillar tau and mature tangle pathology, aged *MAPT* KI P301S mice demonstrate many of the hallmarks of an end-stage human tauopathy. Extending to additional *MAPT* mutations and integrating human biomarker and genetic data, will be essential for defining the full spectrum of tau pathogenic mechanisms.

In summary, this work highlights the heterogeneity of tau pathobiology, identifying mechanisms, timelines and functional vulnerabilities, providing a framework for mechanistic patient stratification and precision medicine in FTD and AD.

## Acknowledgments

We thank Professor Jonathan Rohrer (University College London) for his advice on clinical aspects of human *MAPT* mutations, Professor Malcolm MacLeod (University of Edinburgh) for his help and support on data integrity, Professor Dario Alessi (University of Dundee) for his advice and expertise on proteomic analyses. We thank Dr Marc Diamond for P301S biosensor assay components, Dr Cristina D’Abramo (from the late Peter Davies’ laboratory at Northwell Health) for providing tau antibodies CP13, PHF-1 and MC1, and Dr Nicholas Kanaan (and the late Lester Binder lab at Michigan State University) for TOC1. We also thank Phillip Muckett and Dr Nicholas Cade for technical assistance at the UK DRI at UCL.

## Author contributions

Conceptualization: MF, KED

Methodology: MF, RSN, NW, SB, ET, MB, KED

Investigation: MF, RSN, NW, SB, ET, AS, SP, HD, CG, NF, TB, EC, MW, PMC, IR, DG, MBH, MB

Visualization: MF, MB, DG,

Funding acquisition: MF, KED

Project administration: AAB

Supervision: KED

Writing – original draft: MF, KED

Writing – review & editing: RSN, NW, SB, ET, AS, CG, NF, GB, HZ, NAR, DG, RF, MB

## Data and materials availability

Whole-genome resequencing data from *MAPT* KI, S305N and P301S lines has been deposited in SRA (NCBI) with accession number: PRJNA1152251.

The following datasets were generated for this study: three TMT-phospho-proteomics datasets, DIA phospho-proteomics datasets, snRNA-sequencing dataset, ATAC-sequencing dataset. The datasets generated during and/or analyzed during the current study will be made available from the source data or made publicly available upon publication. All coding performed is available on github: omics analysis, https://github.com/MathieuBo/MAPTKI_proteomic, brain heatmaps https://github.com/MathieuBo/bg-heatmaps and snRNA sequencing analysis https://github.com/eturkes/tau-mutant-snRNAseq.

## Materials and Methods

### Animals

All animal experiments were conducted in accordance with the guidelines of the UK Animal Act 1986 and the Animal Research: Reporting of *In Vivo* Experiments guidelines and were performed at University College London (UCL) under approved UK Home Office project license (PP7490525). C57BL/6J and ICR (Jcl) mice were used as zygote donors and foster mothers. C57BL/6J mice were also used for backcrossing with *MAPT* mutants. *MAPT* KI mice were generated as previously described^22–24^. Preparation of Base Editor, sgRNAs and mouse zygotes for the generation of the *MAPT*^S305N;Int10+3^ KI and *MAPT*^P301S;Int10+3^ KI mouse models has been previously described^24^. APP^NL-G-F^ ^29^ mice were provided by Takaomi Saido, RIKEN, Japan. The *MAPT* KI mice described are available from RIKEN BRC (https://web.brc.riken.jp/en).

### Generation of mutant *MAPT* mutant knock-in mice and off-target assessment

Detailed description of the generation of the *MAPT* mutant knock-in mouse lines has been previously reported^26^. Briefly, third-generation base editing (BE3) was used to generate *MAPT* knock-in (KI) mouse models carrying FTD-associated mutations, consisting of Cas9 nickase (nCas9) fused to rat APOBEC1 and uracil glycosylase inhibitor (UGI). Single-guide RNAs (sgRNAs) were designed to target cytosines within the editable window (positions 4–8) of protospacers adjacent to either NGG (BE3) or NGA (VQR-BE3) PAMs. Two editing sites were selected: the *MAPT*-P301 codon (CCG) and the adjacent intronic splice site *MAPT*-Int10+3. Co-injection of BE3, VQR-BE, and sgRNAs into zygotes derived from heterozygous *MAPT* KI mice resulted in the generation of seven distinct mutant lines. Base conversion at the fourth cytosine of the sgRNA targeting *MAPT*-P301 occurred more efficiently than at the fifth position, leading to a higher frequency of P301L (TTG, 47.6%) compared to P301S (TCG, 11.4%) alleles. Double mutants carrying P301L-Int10+3G>A and P301S-Int10+3G>A were obtained at lower frequencies (5.4% and 2.4%, respectively). All *MAPT*-P301-edited mice also carried a silent V300V (GTC to GTT) mutation. Additional mutant lines included *MAPT*-Int10+3G>A-only mice (21.7%), and rare off-target products such as *MAPT*-P301V (0.6%) and *MAPT*-S305N-Int10+3G>A (0.6%). While multiple mutant *MAPT* mouse lines were established, this study focuses specifically on P301S-Int10+3G>A, *MAPT*-S305N-Int10+3G>A and the non-mutated control *MAPT* KI, aiming to investigate a mutation-dependent phenotype (**Fig. 1A**). Potential off-target sites of BEs in the genome of *MAPT* KI containing S305N and IVS10+3 and P301S and IVS10+3 mutations were assessed using COSMID system (https://crispr.bme.gatech.edu/)^50^. Investigation and assessment of off-target sites for the *MAPT* S305N;Intron10+3 line were described in Watamura et al.^26^. For the *MAPT* P301S;Intron10+3 line, potential off-target sites that accepted up to three mismatches and two insertions and deletions were determined (**Table S1; Fig. S7**) and whole-genome resequencing showed actual off-target mutations (https://www.ncbi.nlm.nih.gov/sra/PRJNA1152251). All sites were removed by backcrossing five times (data was deposited at NCBI with accession number: PRJNA1152251). To assess the off-target mutations, target sites were amplified from tail genomic DNA by PCR using the Ex Taq Polymerase kit (Takara), with primers listed in **Table S2**. Target sequencing was performed using a DNA sequencer (ABI 3730xl). For ease of description, the mutant *MAPT*^S305N;Int10+3^ KI and *MAPT*^P301S;Int10+3^ KI lines will be referred to as S305N and P301S, respectively (**Fig. 1A**).

### Whole Genome Resequencing

#### Library preparation

Genomic DNA was extracted from mouse tail tissue and processed following the Illumina TruSeq DNA sample preparation protocol, to obtain a library of 300-400bp average size. A total of 100 ng of DNA was fragmented using the Covaris system in specialized tubes to produce double-stranded DNA fragments with 3’ or 5’ overhangs. The fragmented DNA was then subjected to end repair to generate blunt ends: 3’ overhangs were removed by exonuclease activity, and 5’ overhangs were filled in by DNA polymerase. Size selection was performed using specific ratios of sample purification beads to isolate fragments of the desired length. A-tailing was then carried out to add a single adenine (A) base to the 3’ ends, allowing for efficient ligation of adapters with a complementary 3’ thymine (T) overhang. Indexed adapters were ligated to the DNA fragments to facilitate multiplexing and subsequent hybridization to the flow cell. The final libraries were amplified by PCR using primers that anneal to the adapter sequences.

#### Clustering and sequencing

Prepared libraries were loaded onto an Illumina flow cell, where DNA fragments hybridized to surface-bound oligonucleotides complementary to the adapter sequences. Clonal clusters were generated via bridge amplification. Sequencing was performed using Illumina’s sequencing-by-synthesis (SBS) technology, which employs reversible terminator-bound dNTPs to enable real-time detection of single base incorporations. This method minimizes incorporation bias and sequencing errors, ensuring high accuracy even in regions with repetitive sequences or homopolymers.

#### Data processing and sequencing

Primary data processing, including image analysis and base calling, was carried out using Illumina’s Real-Time Analysis (RTA) software. The resulting BCL (base call) files were converted to FASTQ format using bcl2fastq2 (version 2.20.0). Sequencing reads were aligned to the reference genome using Isaac Genome Alignment Software (version 01.15.02.08), which provides high accuracy with minimal error rates. Variant detection, including single nucleotide variants (SNVs) and small insertions and deletions (indels), was performed using the Isaac Variant Caller (IVC) (version 2.0.13)^51^. Variant effects on gene function were annotated using SnpEff, which classifies variants based on their predicted impact^52^.

### Antibodies

Antibodies and specific dilutions used in this research project are listed in **Table S3.**

### RNA extraction and real-time PCR on 3R and 4R tau

Total RNA was extracted from mouse brain cortices using RNAiso Plus (Takara) according to the manufacturer’s instructions. Reverse transcription was performed using ReverTra Ace (TOYOBO FSQ-301). Semiquantitative real-time PCR was conducted using the QuantStudio system (Thermo Fisher Scientific). Primer pairs are listed in **Table S4**. The specificity of tau primers was confirmed using the *Mapt* KO mouse.

### RNA extraction and 3R and 4R PCR

RNA was extracted from 10-40mg mouse anterior cortex using the QIAGEN RNeasy Lipid Tissue Mini Kit 74804). Tissue was homogenized with a handheld homogenizer in 1.5mL microcentrifuge tubes with 300uL QIAzol before topping up to 1mL with further QIAzol. Protocol was then followed according to the manufacturer’s recommendations and RNA was eluted in 60uL nuclease-free water, followed by Nanodrop analysis. Fifty ng RNA per sample were used for reverse transcription with NEB LunaScript RT Supermix in 10uL reactions. Four uL of 10x diluted cDNA were used in 10uL PCR reactions using NEB Q5 High-Fidelity 2X Master Mix and custom primers (listed in **Table S5**). PCR cycling conditions were as follows: 95°C for 30s; then 25 cycles of 30s 95°C, 30s 62°C, 60s 72°C; then final extension at 72°C for 120s. One uL of each PCR reaction was analyzed on an Agilent TapeStation 4200 system using a DNA D1000 kit. Molarities for ∼270bp (3R) and ∼370bp (4R) peaks were calculated using the measured ng/uL concentrations and known amplicon sizes (3R: 282bp; 4R: 375bp), rather than using the TapeStation’s calculated peak molarities. Percentage of 4R molarity against total 3R+4R molarity was calculated for each sample. Finally, due to observations of PCR bias, %4R values were adjusted against a standard curve generated by identically amplifying and analyzing synthetic samples consisting of 3R amplicon and 4R amplicon IDT gBlocks mixed at specific ratio increments, for a total DNA concentration of 40fM. Values were interpolated using a cubic curve fit (adj. R^2^ = 0.9971).

### LC-MS/MS analysis targeting tau peptides

Following the protocol described by Lantero-Rodriguez et al.^53^, all brain tissues were homogenized in tris-buffered saline containing a protease inhibitor cocktail (Sigma-Aldrich) and centrifuged at 31,000 × g for 1 hour at 4°C. Protein concentrations were determined from the resulting supernatants. For tau immunoprecipitation, 4 μg of mid-region tau antibody HT7 was conjugated to 50 μL of M-280 Dynabeads (Invitrogen) according to the manufacturer’s instructions. HT7-coated beads were used to immunoprecipitate 10 μg of total protein from brain extracts, adjusted to a final volume of 1 mL with PBS containing 0.05% Triton X-100. To each sample, 10 μL of protein standards mix comprising ¹⁵N-labeled full-length tau was added. To ensure coverage of all tau isoforms, a mixture of 0N3R, 1N4R, and 2N4R isoforms (133 fmol each) was included. Samples were incubated overnight at 4°C on a rolling platform. After washing the beads, tau was eluted with 100 μL of 0.5% formic acid, and eluates were dried under vacuum.

Phosphorylated synthetic peptides labeled with stable isotopes at lysine or arginine residues (Thermo Fisher Scientific) were used as internal standards. The peptides were diluted and mixed, with 100 fmol added to each sample prior to trypsin digestion. Digestion was performed by adding 100 ng of trypsin (Promega) in 50 mM ammonium bicarbonate, adjusting the volume to 50 μL, and incubating overnight at 37°C with 100 rpm rotation. The reaction was quenched with 10% formic acid, and samples were dried in a vacuum centrifuge.

For nanoflow LC-MS analysis, samples were reconstituted in 7 μL of 8% acetonitrile / 8% formic acid. A Dionex 3000 LC system (Thermo Fisher Scientific) coupled to a Q Exactive hybrid quadrupole-orbitrap mass spectrometer (Thermo Fisher Scientific) was used. Six microliters of each sample were loaded onto a reverse-phase Acclaim PepMap C18 trap column (Thermo Fisher Scientific) for desalting and cleanup. Separation was performed on an Acclaim PepMap C18 analytical column (Thermo Fisher Scientific) using a 50-minute linear gradient from 3% to 40% buffer B at 300 nL/min (buffer A: 0.1% formic acid; buffer B: 84% acetonitrile/0.1% formic acid). The mass spectrometer operated in positive ion mode with data-dependent acquisition, acquiring up to five MS/MS scans following each full mass scan. Fragmentation was conducted by higher-energy collisional dissociation (HCD) with normalized collision energy set at 25. Full scan and MS/MS acquisition parameters were set at a resolution of 70,000, 1 microscan, target value of 10⁶, and 250 ms injection time.

LC-MS data were searched against a custom tau database using Mascot Daemon v2.6 and Mascot Distiller v2.6.3 (Matrix Science), with prior charge and isotope deconvolution. Quantitative analysis was performed with Skyline v20.1.0.31 (MacCoss lab) on full scan LC traces. Peaks were manually reviewed and adjusted as necessary, with peptide quantification calculated by the ratio of endogenous to heavy-labeled standard peaks.

### Total-tau ELISA

ELISA INNOTEST® hTAU Ag (Fujirebio) was performed according to the manufacturer’s instructions. Mouse brain extracts were diluted 1:8000 in serial dilutions. Samples were run in singlicate, with one duplicate per group included at the end of each plate. This approach was necessary because the total of 43 samples (three replicates per group) could not be accommodated in duplicate on a single plate. As a quality control, the coefficient of variation (CV%) for the run duplicates was consistently below 10%, and in most cases below 5%.

### Alkaline phosphatase treatment western blot

Mouse brains were homogenized in lysis buffer (50 mM Tris, pH 7.6; 0.15 M NaCl; cOmplete protease inhibitor cocktail) using a Multi-bead Shocker MB (Yasui-Kikai). Homogenates were rotated at 4°C for 1 hour and then centrifuged at 19,800 × g for 30 minutes. Supernatants were collected as lysates and subjected to SDS-PAGE, followed by transfer onto PVDF membranes. To assess tau isoform patterns, brain lysates were dephosphorylated with lambda phosphatase (Santa Cruz Biotechnology) according to the manufacturer’s instructions. Membranes were blocked with ECL Prime blocking buffer (GE Healthcare) for 1 hour and incubated overnight at 4°C with primary antibodies (**Table S3**). Immunoreactive bands were detected using ECL Select detection reagent (GE Healthcare) and imaged on a LAS-3000 Mini Lumino image analyzer (Fujifilm). Quantification was performed using ImageJ (v1.53a) and Fiji software (NIH).

#### Total and phosphorylation proteomics

Phosphopeptide enrichment and TMT-labelling was performed according to accessible protocol^54^, as previously published^55^.

#### Tissue Sample Processing

Frozen mouse hippocampi were weighed and processed on dry ice. Tissue was homogenized using a CryoLys Evolution homogenizer (3 cycles at 2000 rpm) in 1 mL SDS lysis buffer (2% SDS in 100 mM TEABC, pH 8.5, with protease and phosphatase inhibitors). Samples were centrifuged at 2,000 × g for 2 min, transferred to low-bind tubes, boiled at 95°C for 5 min, sonicated (15 cycles of 30 s on/off), and centrifuged at 20,000 × g for 20 min. Protein quantification was performed on the supernatants by Pierce BCA Protein Assay kit.

#### S-Trap-Assisted Protein Digestion

Protein (3 mg) was reduced with 0.1 M TCEP (10 mM final) at 60°C for 30 min, alkylated with 40 mM M iodoacetamide at room temperature in the dark, and quenched with TCEP. The final SDS concentration was adjusted to 5%. Samples were diluted 7-fold with S-Trap binding buffer, loaded onto S-Trap midi columns, centrifuged, washed, and digested with trypsin/Lys-C (47°C, 1.5 h, then overnight at room temperature). Peptides were eluted with TEABC, 0.15% formic acid, and 50% acetonitrile, pooled, dried, and stored at -80°C.

Sep-Pak Purification: Peptides were reconstituted in 1% TFA, incubated at room temperature (30 min, 1800 rpm), centrifuged (17,000 × g, 10 min), and purified using Sep-Pak tC18 cartridges. Columns were activated, equilibrated, loaded with peptides, washed with 0.1% formic acid, and eluted with 50% acetonitrile in 0.1% formic acid. At this point, 5% of eluate was aliquoted separately for total proteome analysis. Dried eluates were stored at -80°C.

#### Phosphopeptide Enrichment Using TiO_2_

Peptides were mixed with binding buffer, centrifuged, and loaded onto TiO_2_ spin tips (Thermo Fisher Scientific). After washing, phosphopeptides were eluted, pooled, centrifuged (17,000 × g, 30 s), snap-frozen, dried, and stored at -80°C.

TMT Labelling of Peptides: Peptides were dissolved in 50 mM TEABC and incubated (20 min). TMT reagents were added and incubated for 2 h, followed by quenching with 5% hydroxylamine (10 min). Labelled peptides were pooled, dried, and submitted for high-pH fractionation.

#### LC-MS/MS Analysis (TMT)

Peptide fractions were reconstituted in 10 mM ammonium formate (pH 10.0) and analyzed using a Dionex RSLC 3000 nano-LC and Orbitrap Fusion Lumos mass spectrometer. Analysis included both phosphoproteomic and total proteome runs. MS data were searched using MaxQuant or MS-Fragger, and results were analyzed with Perseus.

#### LC-MS/MS analysis (DIA)

DIA-MS for phosphoproteome and total proteome is acquired on an Orbitrap Astral mass spectrometer interfaced with Vanquish-Neo liquid chromatography system. Phosphopeptides were dissolved in LC buffer (0.015% (v/v) DDM, 3% ACN (v/v), 0.15% formic acid in LC-water). Peptide were loaded using trap and elute and resolved on Aurora Ultimate™ 25cm ×75 µm XS C18, 1.7µm pore size UHPLC column (IonOptiks). The LC and mass spectrometer was operated for 30 min to achieve 40 samples per day throughput. Full MS scan was acquired using Orbitrap mass analyzer at a resolution of 180, in the scan range of 420-1020 m/z. DIA scans were acquired using narrow-window isolation scheme set to 4 m/z with a total of 150 scans were acquired per MS2 cycle. Precursor ions were accumulated with a AGC target of 500% for 3.5ms. Precursor ions were fragmented 27% HCD and measured using Astral mass analyzer. Total proteome data was acquired using the above parameters except the MS1 scans acquired at a resolution of 240,000 and measured using Orbitrap mass analyzer. DIA scans acquired in the scan range of 380-980 with a isolation window set to 2 m/z comprising of 299 MS2 scans and precursor ions were fragmented with 25% HCD and measured using Astral analyzer (Thermo Fisher Scientific).

#### Data Analysis (TMT)

For the TMT-phospho-proteomics datasets (see specifics in figure legend) several analysis methods were applied. Firstly, since protein function is modulated by site-specific phosphorylation or cumulative phosphorylation of multiple phosphosites, we measured the overall phosphorylation status (ΔPs) of phosphoproteins in our dataset, as the sum of log2(fold change) value of all phosphopeptides with statistically significant changes (p < 0.05) compared to control. Tau phospho-peptides were normalized by total-tau peptides. If none of the phosphopeptide P values is below 0.05, the ΔPs value will be zero. A stringent cut-off for ΔPs value was applied at three standard deviations (3σ), which revealed hyperphosphorylation (ΔPs > 3σ) and hypophosphorylation (ΔPs < −3σ). Secondly, eigenprotein scores were calculated by performing principal component analysis (PCA) on z-score normalized intensity values of phospho-tau peptides across samples. The first principal component (PC1), representing the dominant axis of variation, was extracted and used as the eigenprotein score for each sample. Heatmaps comparing the abundance of specific phospho-tau peptides were produced to visualize the behavior in the different mouse lines.

#### Data Analysis (DIA)

The raw DIA MS data from Phosphoproteome and total proteome sets have been processed using Spectronaut search engine V20.1 (Biognosys). The data was searched against a mouse UniProt database by including human Tau protein (441 aa) of wild-type knock-in and mutant versions. The search parameters for total proteome includes Trypsin as a strict protease; 2 missed cleavages allowed; minimum peptide length was set to 7 amino acids; Carbamidomethylation of Cys as fixed and phosphorylation of STY, Oxidation of Met, Deamidation of NQ and protein-N-termini acetylation as variable modifications. PSM, peptide and protein-group level FDR was set to 1%; precursor q-value of 0.01 and precursor PEP cutoff set to 0.2; direct-DIA search is enabled and PTM analysis was set to default by setting the localization score threshold cut-off set to 0.75; NO imputation and cross-run normalization is disabled, and other parameters were set default. The PTM sites report is exported and further processed using Perseus version 1.6.015 and in-house generated python scripts. The total proteome searches were done using the above and default parameters. Single-hit proteins were excluded for quantification, global imputing and cross-run normalization enabled. The protein group files were exported and further analyzed using Perseus (v1.6.0.15).

#### Data visualization

was performed using Python. For the TMT-total proteome, differentially accumulated proteins were identified and normalized relative to *MAPT* KI, and significantly altered proteins (p-value<0.05) were subjected to Gene Ontology (GO) enrichment analysis using BiNGO within Cytoscape, to map enriched biological processes.

### Western blot

Isopentane frozen and dissected mouse brain hippocampi were homogenized in RIPA lysis & extraction buffer (Thermo Fisher Scientific; with an addition of 1/100 HALT Protease and Phosphatase Inhibitor Cocktail, Thermo Fisher Scientific) using a hand-held automatic homogenizer. Homogenates were left to solubilize for 30 min at 4°C and then centrifuged at 10,000 × g for 10 minutes. Supernatants were collected and protein concentration was measured by BCA Protein Assay kit. Supernatants were subjected to SDS-PAGE on 4-20% Criterion TGX Stain-Free Protein gels (Bio-Rad) and proteins were transferred to low-fluorescence PVDF membranes (BioRad). Following visualization using UV light in a Chemi-Doc MP Imager (Bio-Rad), membranes were blocked with 5% milk in tris-buffered saline (20 mM Tris-HCL pH 7.4, 150 mM NaCl) containing 0.02% Tween-20 (TBST) for 1h at room temperature (RT). Primary antibodies (**Table S3**) were diluted in Superblock TBS blocking buffer (Thermo Fisher Scientific) and incubated overnight at 4°C. Membranes were washed twice in TBST and twice in 5% milk for 10 min each. Secondary HRP-conjugated secondary antibodies were diluted in 5% milk and incubated for 1h at RT. Membranes were washed 4x in TBST for 10 min each and immunoreactive protein bands were detected using Clarity Western ECL substrate (Bio-Rad) and imaged on ChemiDoc MP Imager. Quantification was performed using ImageLab (v6.1.0).

### Immunohistochemistry and immunofluorescence

Mouse brains were perfused with PBS and 4% paraformaldehyde (Merck). For tau immunohistochemistry, post-perfusion, brains were incubated in 30% sucrose in PBS for 48 hours at 4°C and subsequently embedded in OCT compound (Sakura Finetek).

#### Tau immunohistochemistry staining

Free-floating horizontal sections (30 μm) were rinsed in PBS and treated with 0.3% H_2_O_2_ in methanol for 15 minutes at RT to quench endogenous peroxidase activity. Sections were then blocked for 1.5 hours in M.O.M. blocking solution (Vector Laboratories) and incubated overnight at 4°C with primary antibodies (listed in **Table S3**) diluted in the reagent provided by the M.O.M kit. After washing in PBS, sections were incubated with secondary antibodies for 15 minutes, followed by signal amplification using the avidin-biotin complex (ABC, Vector Laboratories) for 10 minutes. Immunoreactivity was visualized using 3,3′-diaminobenzidine (DAB) with H_2_O_2_ activation. Sections from seven to ten mice per group were processed simultaneously in a genotype-blinded manner. A semiquantitative scoring system based on signal intensity and extent of labeling (0: none; 1: mild; 1.5: moderate; 2: severe) was used to assess staining intensity across defined anatomical regions, independently rated by two blinded investigators. Representative anatomical heatmap illustrations were created with BrainGlobe-Heatmap^56^. For each brain, pathology was assessed across one bregma (coronal bregma -3.15mm from Paxions and Franklin’s mouse atlas) in several Ns (see legend for details).

#### Immunofluorescence staining

Formalin-fixed paraffin-embedded 8 µm sections were deparaffinized in xylene and rehydrated through a graded ethanol series. Antigen retrieval was performed by pressure cooker treatment in citrate buffer for 10 minutes. Sections were then blocked in M.O.M. blocking buffer and incubated with primary antibodies (diluted in M.O.M. diluent) (**Table S3**) overnight at 4°C. After washing, sections were incubated in Alexa Fluor (Thermo Fisher Scientific) secondary antibodies, followed by a 10 min DAPI incubation. Sections were mounted in ProLong Antifade Mounting medium (Invitrogen).

### Cell Culture and Seeding Assay

HEK293 Tau RD P301S FRET biosensor cells (ATCC CRL-3275) and HEK Tau RD S305N-YFP biosensor cells were maintained in DMEM (Thermo Fisher Scientific) supplemented with 10% fetal bovine serum and 1% penicillin-streptomycin. Cells were seeded at a density of 35,000 cells per well in 96-well plates, with a total volume of 130 µL per well. For imaging experiments, cells were plated in poly-L-ornithine-coated PhenoPlate 96-well microplates (Perkin Elmer). After 18 hours, cells were transduced with proteopathic tau seeds. Briefly, 1.25 µL of Lipofectamine 2000 (Invitrogen) was mixed with 8.75 µL of Opti-MEM (Gibco) and incubated at room temperature for 5 minutes before combining with 10 µg of total protein from mouse brain extracts. Negative control cells received Lipofectamine in Opti-MEM without seed material. After a 30-minute incubation at 37 °C, 20 µL of the transfection mix was added to each well. Cells were incubated for 48h prior to analysis. All conditions were tested in triplicate.

#### Flow Cytometry Analysis for P301S biosensor line

To quantify tau seeding, HEK Tau RD P301S FRET biosensor cells were harvested using 0.25% trypsin, fixed in 4% paraformaldehyde for 10 minutes, and resuspended in flow cytometry buffer (1X PBS with 1 mM EDTA). FRET flow cytometry was performed using an LSR Fortessa Flow Cytometer (BD Biosciences). FRET-positive cells were identified as previously described. In brief, single cells double-positive for CFP and YFP were gated, and FRET-positive events were quantified within this population. Percentage of FRET-positive cells was calculated as output metric. Data were analyzed using FCS Express v7 (De Novo Software) and GraphPad Prism.

#### Imaging and spot count analysis for S305N biosensor line

Tau RD S305N-YFP biosensors were fixed in 4% PFA for 10 min, washed three times in DPBS, and stained with Hoechst (Invitrogen, cat. no. 33342, 1:1,000) for 10 min at room temperature. Following three additional DPBS washes, cells were stored in PBS at 4 °C until imaging on the Opera Phenix Plus high-content confocal microscope (PerkinElmer). Excitation and emission settings were as follows: YFP, 488 nm excitation with 650–760 nm emission; Hoechst, 375 nm excitation with 435–480 nm emission. Images were acquired using a ×20 water-immersion objective, with 12 fields of view per well and a Z-stack of seven slices at 1.5 μm intervals. Image analysis was performed in Columbus software v.2.9.1 (PerkinElmer). Maximum projections were generated and basic flatfield correction was applied. Nuclei were segmented using the Hoechst channel, and cytoplasm was subsequently identified by watershed expansion from nuclei. YFP-positive tau inclusions were detected using the ‘find spots’ building block (method D) on the YFP channel. The number of spots per nucleus was calculated from exported data.

#### 4R tau Real-Time Quaking-Induced Conversion (RT-QuIC) Assay

RT-QuIC was carried out as previously described^28^. Briefly, the recombinant tau protein K11CFh—a fragment encompassing the R1–R4 domains up to residue 400 with cysteine-to-serine substitutions—was purified and used to amplify tau seeds from 10% (w/v) PBS (with protease and phosphatase inhibitors cocktail) brain extracts. These extracts were derived from the frontal cortex of human PSP cases and from the posterior cortex of 15M *MAPT* KI, S305N and P301S mice. K11CFh was thawed from −80 °C and passed through 100 kDa molecular weight cut-off filters to eliminate preformed aggregates. RT-QuIC reactions were seeded with a 1 × 10⁻⁵ dilution of brain homogenate in the presence of 3 µM K11CFh and 10 µM thioflavin T (ThT) in reaction buffer (250 mM trisodium citrate and 10 mM HEPES, pH 7.4). Reactions were set up in 384-well Nunc microplates and sealed with an aluminum cover to prevent evaporation. Plates were incubated at 37 °C in a BMG FluoStar Lite plate reader using alternating cycles of 60 seconds of orbital shaking at 500 rpm and 60 seconds of rest. ThT fluorescence was recorded every 15 minutes to monitor aggregation kinetics.

### Sarkosyl-extraction

Whole brains (without cerebellum and olfactory bulb) were homogenized in 10% extraction buffer (10mM Tris, pH 7.4, 0.8M NaCl, 1mM EGTA, 5mM EDTA, 10% sucrose) with HALT Protease and Phosphatase Inhibitor Cocktail (Thermo Fisher Scientific) using a handheld homogenizer. Homogenates were left to solubilize for 30 min on ice and centrifuged at 10,000 × g for 10 minutes at 4 °C. Supernatants were collected, and the remaining pellet was sequentially re-extracted three additional times. Protein concentration was measured using the Pierce BCA Protein Assay kit. Equal total protein volumes were calculated, and each supernatant was treated with sarkosyl to a final concentration of 1% and incubated at room temperature for 1 hour with gentle rotation. Samples were then centrifuged at 300,000 × g for 70 minutes. The resulting pellets, containing sarkosyl-insoluble tau, were washed by resuspension in 50 mM Tris-Hcl pH 7.4 containing 150 mM NaCl and re-spun at 300,000 × g for 70 minutes. The final pellet was resuspended in 10µL of 50mM Tris-HCL pH 7.4. The same protocol was used to produce samples for immunogold labelling and negative stain (below).

### Immunogold labelling

Sarkosyl-insoluble fractions were prepared from whole brains as described in “sarkosyl-extraction”, and resuspended in PBS. 100 mesh nickel formvar/carbon coated grids were glow discharged in a PELCO easiGlow glow discharging system for 30 seconds. 5ul of resuspended sarkosyl-insoluble solution was placed on the grid and incubated for 5 minutes. The grid was then rinsed three times with 100ul of PBS for 10 minutes each time. Grids were then treated with 50ul of 0.05M glycine for 10 minutes before being transferred to a drop of blocking buffer for 30 minutes. Grids were then incubated with 10ul of primary antibody (MC1 or HT7) at a dilution of 1:2 overnight at 4C. The next day the grid is washed with five separate drops of PBS with 0.1%BSA for 10 minutes each. The grid is then transferred to the secondary antibody for 1 hour (Secondary antibody conjugated to 6nm gold and used at 1:50). The grid is subsequently washed five times in PBS with 0.1%BSA for 10 minutes each and two washes in distilled water. Negative staining is then performed by incubating the grid with 2% Uranyl acetate for 5 minutes. Excess stain is removed by using filter paper and then is imaged in a Talos L120C transmission electron microscope operating at 120kV. Images were taken using a Ceta camera.

### Nuclei isolation and single-nucleus RNA sequencing

Nuclei isolation was performed as previously described^57^. Briefly, hippocampi from 24-month-old mice (N=4 per genotype; *MAPT* KI, S305N, and P301S) were dissected, snap frozen, and processed for nuclei isolation. Frozen tissue was sliced on dry ice and homogenized using a glass Dounce homogenizer (15 gentle strokes) in 1 mL ice-cold Homogenization Buffer (HB: 320 mM sucrose, 5 mM CaCl₂, 3 mM Mg-acetate, 10 mM Tris, 0.1 mM EDTA, 0.1% Igepal, 0.1 mM PMSF, 1 mM 2-mercaptoethanol, RNasin Plus). The homogenate was passed through a 70 μm strainer and washed with 1.65 mL HB to a final volume of 2.65 mL. An equal volume (2.65 mL) of Gradient Medium (50% Optiprep, 5 mM CaCl₂, 3 mM Mg-acetate, 10 mM Tris, 0.1 mM PMSF, 1 mM 2-mercaptoethanol) was added (final volume 5.3 mL). Samples were then layered onto a 4 mL 29% Optiprep cushion prepared in Optiprep Diluent Medium (ODM: 150 mM KCl, 30 mM MgCl₂, 60 mM Tris, 250 mM sucrose) and centrifuged at 7,700 rpm for 30 min at 4 °C in a SW41Ti rotor. The supernatant was removed, and the nuclei pellet was gently resuspended in 200 μL resuspension buffer (1× PBS, 1% BSA, 0.2 U/mL RNasin Plus). The suspension was washed with an additional 100–200 μL resuspension buffer, pooled, and filtered through a 35 μm strainer to remove clumps. Nuclei yield, integrity, and morphology were assessed using C-Chip hemocytometers. In addition, 18µl of sample and 2µl of Acridine Orange/Propidium iodine were added and loaded onto a LUNA-FL slide. cDNA libraries generated from fresh-frozen nuclei were sequenced on an Illumina NovaSeq X Plus on a 10B flow cell using 300-cycle kit with 15% phiX spike-in, with a target of 30,000 reads per nucleus. Reads were demultiplexed to FASTQ files using bcl2fastq and then mapped to the mouse reference genome (mm10).

SnRNA-seq libraries were generated on the 10x Genomics Chromium platform and processed with Cell Ranger with the inclusion of intronic data^58^. Analyses began from the unfiltered matrix. For each sample, count data were imported into R and assembled as a Seurat object^59–62^. Gene annotation was taken from the accompanying features file to map Ensembl identifiers to gene symbols. To characterize ambient RNA and examine barcode rank behavior, we used the R package DropletUtils^63,64^. Barcode rank curves were inspected at the knee and inflection points as diagnostics. Cell-containing droplets were then called with emptyDrops, and only barcodes with a false discovery rate at or below 0.001 were retained for downstream analysis.

Standard per-nucleus quality metrics were computed in Seurat^65,66^. Mitochondrial genes were used to derive the mitochondrial fraction per nucleus, ribosomal protein genes to derive the ribosomal fraction, and *malat1* to summarize nuclear RNA content. Using the estimated ambient RNA profile from DropletUtils, we also assessed potential contamination trends. Because the data consisted of single-nuclei rather than whole-cell libraries, mitochondrial-encoded genes were removed from the count matrix before normalization.

Putative doublets were then identified with scDblFinder^67^. scDblFinder operates under the assumption that empty droplets have been removed, but further QC had not been performed. Therefore, called doublets were removed prior to nuclei removal based on the QC metrics previously described. After assessing these metrics, we applied a conventional thresholding of nuclei with mitochondrial fraction of 10% or greater^66^. We also plotted density estimations of unique features and total RNA counts for each nuclei, setting a fine-tuned threshold for each sample to remove the large density of “low-quality” nuclei consistently observed. We determined this threshold to be around 200 detected genes and 200 total counts for most samples.

Following these steps, the samples were combined into a single Seurat object. Normalization, dimensionality reduction, and clustering followed standard Seurat protocol, using the SCTransform^68^, UMAP^69^, and Seurat louvain clustering methods, respectively. Principal component selection was driven by the identification of inflection points on Elbow plots. To aid in the annotation of cell-types, we aligned our data with the Allen Brain Atlas^70^ by using the Seurat reference-query approach. Based on the results of the alignment, we manually annotated the clusters produced from the louvain method first into broad categories (*e.g*. neurons, glial types, etc.). These broad types were then subclustered iteratively using the normalization, dimensionality reduction, and clustering approach described until sufficient resolution was achieved for fine-grained cell-typing.

snRNAseq-based differential expression was performed using a zero-inflated negative binomial model (MAST^71^) in Seurat and FDR correction of the p-values. Differential expression was assessed in both S305N and P301S lines compared to *MAPT* KI at the cell-type level (*i.e*. CA3 neurons). Each list of differentially expressed genes was then independently submitted to BINGO for pathway analysis and integrated in Cytoscape using EnrichmentMap.

### EM analysis of cytoskeleton

Mice were transcardially perfused with a solution of 4% PFA and 2% glutaraldehyde in 0.1 M sodium cacodylate buffer, pH 7.4. Fixed tissue was cut into 50-μm-thick vibratome sections and postfixed in 1% osmium tetroxide for 1 hour, washed and then incubated in 2% uranyl acetate solution at 4C overnight. Following alcohol dehydration, tissue sections were infiltrated with increasing concentrations of Spurr resin and then embedded flat in Aclar sheets. Regions of interest for ultrastructural analyses were excised, and 50 nm ultrathin sections were prepared using an ultramicrotome on to 75 mesh formvar/carbon grids before being stained with 2% uranyl acetate and lead citrate. The thin sections were then viewed using a Thermo Fisher Talos L120C transmission electron microscope operating at 120 kV. Ten images were acquired per biological replicate. Microtubule morphology was quantified from images using Fiji (ImageJ) and analysis were performed blind by three independent researchers. Spatial calibration was performed globally to all images by measuring the scale bar several times and converted based on the pixel to nm ratio. Individual microtubules were identified manually, and their lengths and widths were measured using the straight-line tool and ROI Manager. The number of microtubules per image was determined from the count of individual length measurements. Measurements were exported to Excel, unblinded and grouped by experimental condition for subsequent statistical analysis on GraphPad Prism.

### ATAC-sequencing

Nuclei were isolated from frozen mouse hippocampal tissue as described^57^. Nuclei yield, integrity, and morphology were assessed using C-Chip hemocytometers. In addition, 18µl of sample and 2µl of Acridine Orange/Propidium iodine were added and loaded onto a LUNA-FL slide. Bulk ATAC-seq was carried out as in Corces et al.^72^, using 2µl Tn5 transposase using 35,000 nuclei per transposition reaction. Libraries were examined using Tapestation 4200 using D5000 Screentape and reagents. Bulk ATAC-seq libraries were sequenced using an Illumina NovaSeq-X Plus instrument on a 10B flowcell using a 300-cycle kit with 15% phiX spike-in.

### Negative stain

Sarkosyl-soluble samples (3 µl per grid) were applied to 15 s glow-discharged (Pelco easyglo) continuous carbon grids (Agar scientific) for 60 s, after which grids were dried by wicking the sample solution with filter paper (Whatman #1). EM grids were then washed 3 times by applying and wicking 5 µl of water, stained with 5 µl of 2% w/v uranyl acetate for 30 s, after which grids were dried by wicking the sample solution with filter paper. This staining step was repeated a second time before EM grids were imaged on an ABSL (Leeds) Talos 120 kEV (Thermo Fisher Scientific) or Technai T12 (Thermo Fisher Scientific) transmission electron microscope at 13500x and 92000x magnification.

### Cryo-EM

Plunge frozen sarkosyl-soluble samples for cryoEM were prepared by applying 3 µl of sample to 60 s plasma cleaned (Tergeo, Pie Scientific) Quantifoil R1.2/1.3 grids. Grids were blotted and plunge-frozen in liquid ethane using a Vitrobot Mark IV (Thermo Fisher Scientific) with a 1 s wait and 5 s blot time respectively. The Vitrobot chamber was maintained at close to 100% humidity and 6°C. The single-particle datasets were collected at ABSL (Leeds) using a Titan Krios electron microscope (Thermo Fisher Scientific) operated at 300 ke−V with a Falcon4 detector in counting mode (Nominal magnification of 96,000x and 0.83 Å/pixel). A total of 89 movies were collected using EPU-3.0 (Thermo Fisher Scientific) with a nominal defocus range of −1.6 to −3.1 µm and a total dose of ∼52 e−/Å2 over an exposure of 8 s, corresponded to a dose rate of ∼4.5 e−/pixel/s.

Raw EER movies were compressed and converted to TIFF using RELION-4^73^, regrouping frames into 40 fractions to give a dose per frame of 1.3 e−/Å2. The TIFF stacks were aligned and summed using motion correction in RELION-4 and CTF parameters were estimated for each micrograph using CTFFIND4^73^. Filaments were manually picked and 2D class averages were generated. Fast Fourier transform of summed micrographs was performed in Python.

### Statistics

For comparisons of two groups, statistical analyses were conducted using Welch’s t-test. For comparisons involving three or more groups, one-way or two-way analysis of variance (ANOVA) were conducted, depending on the number of independent variables, followed by *post hoc* testing. When either the normality or equal variance assumptions were not satisfied, data were analyzed with Kruskal–Wallis one-way ANOVA on ranks followed by Dunn’s multiple comparisons test. To assess statistical difference following eigenprotein analysis, one-way ANOVA with Tukey’s Honestly Significant Difference was performed. Specific statistical analysis for each experiment can be found in the statistical summary file. All analyses and visualizations were performed in GraphPad Prism (v.10.6.0) or Python (v3.13.5, Python Software Foundation). Each dot in a barplots indicates biological replicates; for heatmap visualization and western blotting technical replicates are also displayed. For RT-QuIC technical replicates are displayed to show variability at the peak stage reaction. Details on Ns used for each experiment can be found in the figure legend. All presented data are representative of the same experiment performed in at least three animals (except in the NLGF-MAPT KI seeding at 30M due to lack of mice available at such an advanced age). All experiments were replicated in at least two independent experiments unless stated otherwise. No statistical methods were used to pre-determine sample sizes but our sample sizes are similar to those reported in previous publications^23–25,74^. Animals were randomly assigned to experimental groups, and investigators were blinded to genotype throughout the experiments unless blinding was not feasible. Specific details are found within each method. PCA, Eigenprotein analyses, ΔPs, heatmaps were conducted in Python using the following packages: SciPy (v1.8.1), NumPy (v1.23.1), Scikit-learn (v0.19.3), Seaborn (v0.13.2), Matplotlib (v3.5.2), Brainglobe-Heatmap (0.6.0), Statsmodels.api (v0.14.4), Pandas (v2.2.3).

