## supplemental and table material for "Distinct mechanistic pathways of early tauopathy revealed by *MAPT* mutations"

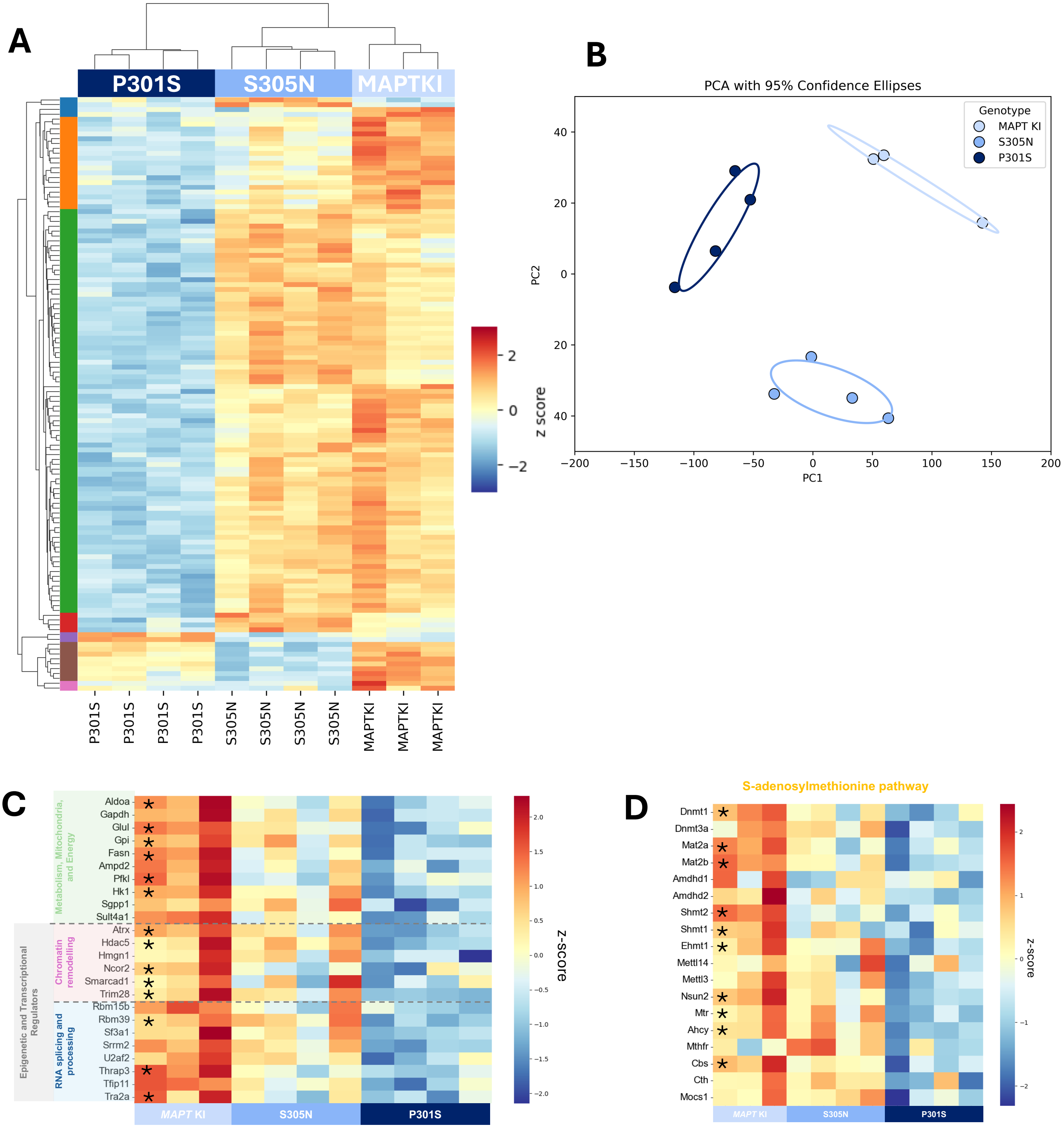
A
P301S
S305N
MAPTKI
z score
B
PCA with 95% Confidence Ellipses
Genotype
MAPT KI
S305N
P301S
PC1
PC2
C
Metabolism, Mitochondria, and Energy
Chromatin remodelling
RNA splicing and processing
Epigenetic and Transcriptional Regulators
Aldoa
Gapdh
Glul
Gpi
Fasn
Ampd2
Pfkl
Hk1
Sgpp1
Sult4a1
Atrx
Hdac5
Hmgn1
Ncor2
Smarcad1
Trim28
Rbm15b
Rbm39
Sf3a1
Srrm2
U2af2
Thrap3
Tfip11
Tra2a
MAPT KI
S305N
P301S
Z-score
D
S-adenosylmethionine pathway
Dnmt1
Dnmt3a
Mat2a
Mat2b
Amdhd1
Amdhd2
Shmt2
Shmt1
Ehmt1
Mettl14
Mettl3
Nsun2
Mtr
Ahcy
Mthfr
Cbs
Cth
Mocs1
MAPT KI
S305N
P301S
Z-score

**Fig. S1.** (**A**) Cluster heatmap of total proteomics dataset comparing 15M posterior cortex from MAPT KI (N=3; N=1 male; N=2 female); S305N (N=4; sex matched) and P301S (N=4; sex matched) and (**B**) PCA showing data distribution across dataset from (A). Ellipses represent 95% confidence intervals (**C**) Heatmap showing protein level abundance of the proteins in the P301S-specific cluster (named metabolism and epigenetic regulators). * indicate proteins that show a $p<0.05$ with FDR correction between *MAPT* KI and P301S. **(D**) Heatmap showing protein level abundance of SAM (S-adenosylmethionine)–pathway. * indicate proteins that show a $p<0.05$ with FDR correction between *MAPT* KI and P301S.

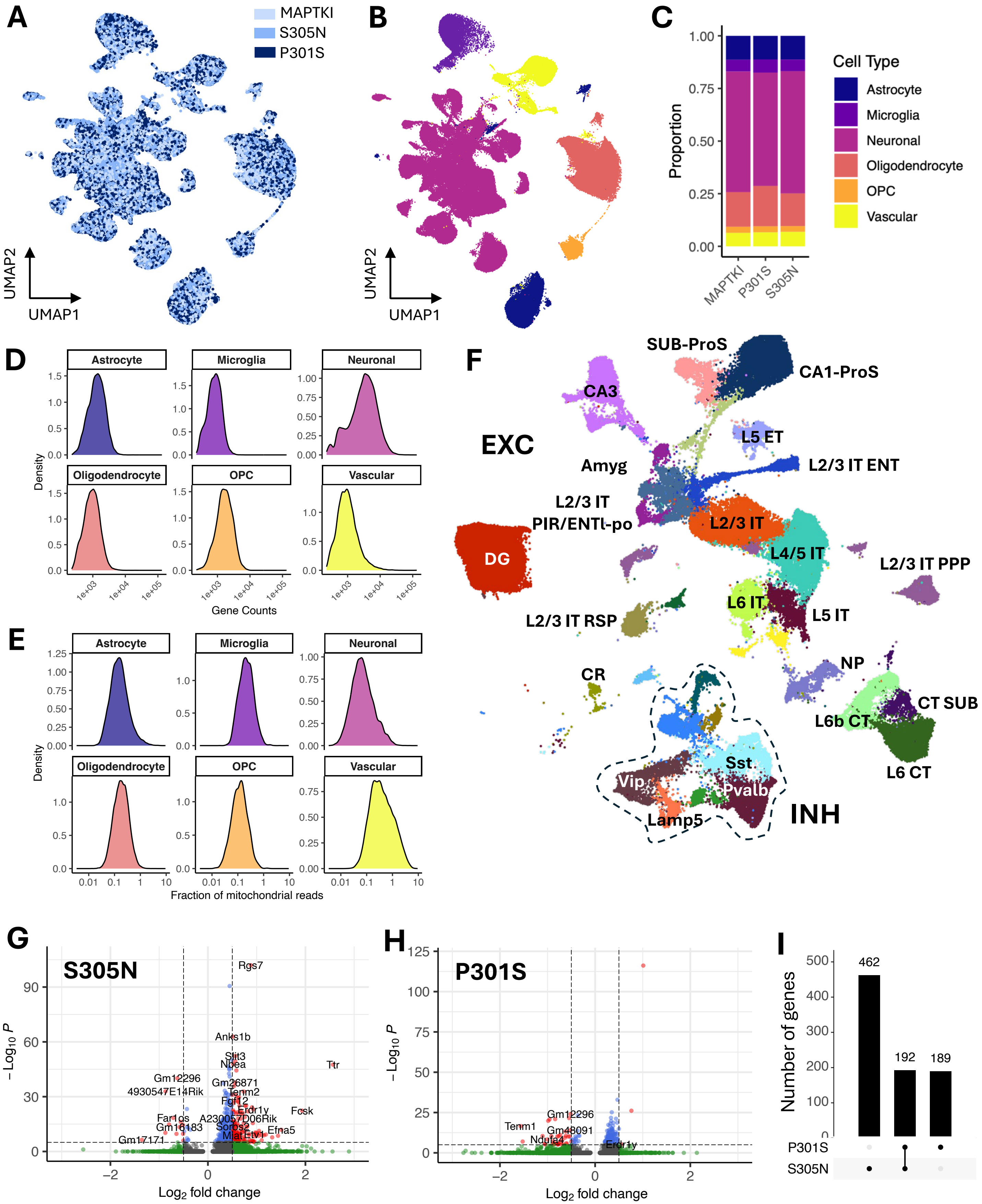

A
MAPTKI
S305N
P301S
UMAP2
UMAP1
B
C
Proportion
Cell Type
Astrocyte
Microglia
Neuronal
Oligodendrocyte
OPC
Vascular
MAPTKI
P301S
S305N
D
Density
Gene Counts
E
Fraction of mitochondrial reads
F
EXC
SUB-ProS
CA1-ProS
CA3
L5 ET
Amyg
L2/3 IT ENT
L2/3 IT
PIR/ENTl-po
L2/3 IT
L4/5 IT
DG
L2/3 IT PPP
L6 IT
L5 IT
L2/3 IT RSP
CR
NP
CT SUB
L6b CT
L6 CT
Sst
Vip
Pvalb
Lamp5
INH
G
S305N
Rgs7
Anks1b
Slit3
Nbea
Ttr
Gm12296
Gm26871
4930547E14Rik
Tenm2
Fgf12
Erdr1y
Fosk
Far1os
A230057D06Rik
Gm16183
Sorbs2
Efna5
Mlat
Etv1
Gm17171
Log2 fold change
H
P301S
Gm12296
Tenm1
Gm48091
Ndufa4
Erdr1y
I
Number of genes
462
192
189
P301S
S305N

**Fig. S2** (**A,B**) UMAP embedding of single nuclei RNA sequencing data from CA3 of hippocampus of 24M *MAPT* KI, S305N and P301S mice (N=4, sex matched). Each dot represents a cell that is coloured according to the genotype (**A**) or broad cell-type annotation (**B**). OPC: oligodendrocyte progenitor cells. (**C)** Relative proportion of broad cell-types between lines (info in A,B). (**D,E**) Density plots per cell-types of gene counts (**D**) and fraction of mitochondrial reads (**E**). Note that x-axes are displayed in log scale. (F) UMAP embedding of neuronal cells. Each cell is coloured according to its fine-grained identity (Supplementary table 6 for a full description). (**G,H**) Volcano plots displaying differentially expressed genes (in red, |Log2FC| > 0.5 and adjusted p-value < 0.05) between S305N and *MAPT* KI (**G**) and P301S and *MAPT* KI (**H**). (**I**) Upset plot highlighting the fraction of shared or unique differentially expressed genes for each comparison.

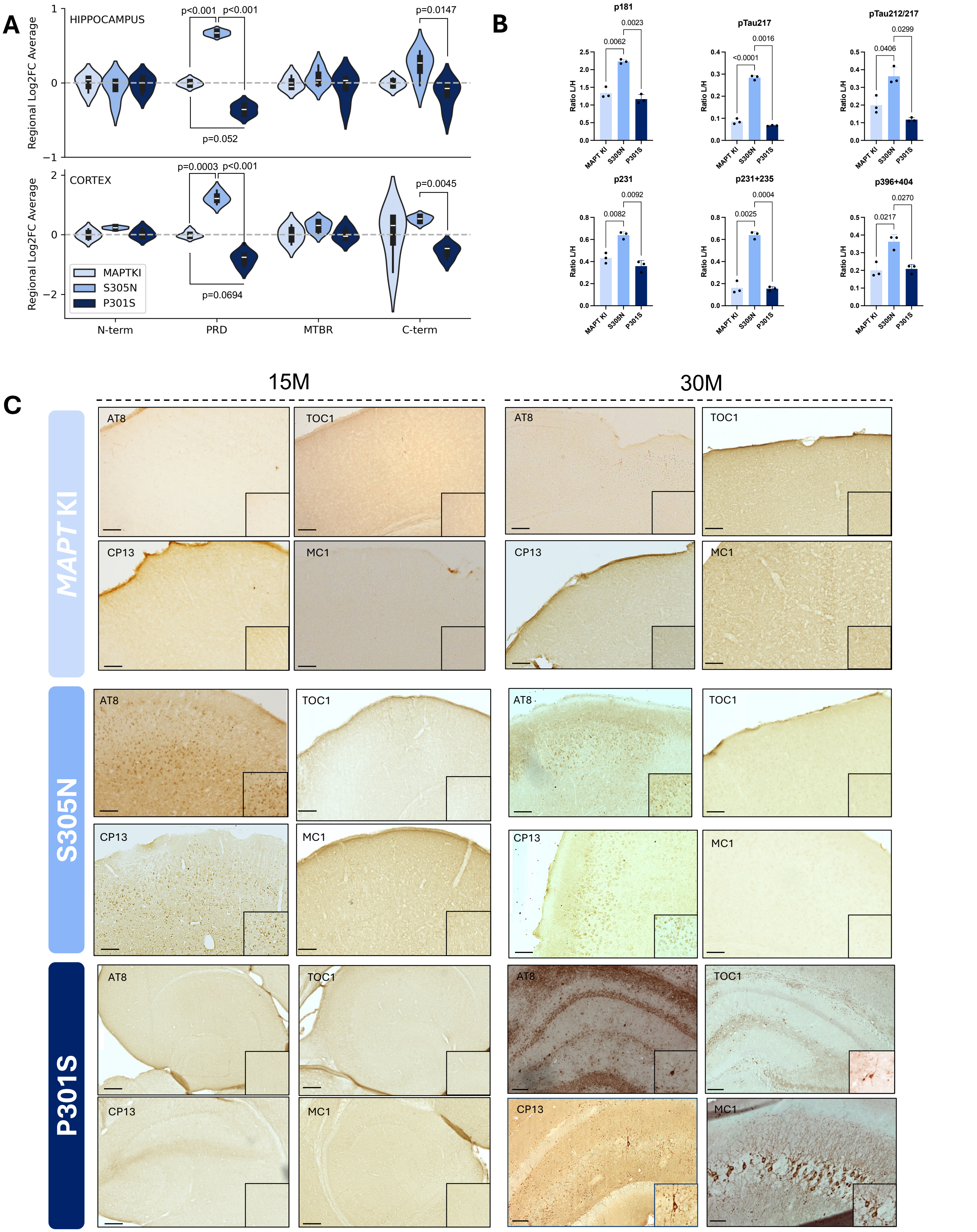
A
HIPPOCAMPUS
Regional Log2FC Average
p<0.001
p<0.001
p=0.0147
p=0.052
CORTEX
p=0.0003
p<0.001
p=0.0045
p=0.0694
MAPTKI
S305N
P301S
N-term
PRD
MTBR
C-term
B
p181
0.0062
0.0023
pTau217
<0.0001
0.0016
pTau212/217
0.0406
0.0299
p231
0.0082
0.0092
p231+235
0.0025
0.0004
p396+404
0.0217
0.0270
Ratio L/H
MAPT KI
S305N
P301S
C
15M
30M
MAPT KI
S305N
P301S
AT8
TOC1
CP13
MC1

**Fig. S3. (A)** Eigenprotein values for phospho-tau–associated peptides were derived from principal component analysis of the TMT-based phospho-proteomics dataset 1 and 3 subdivided by tau regions, *i.e.* N-terminus (aa:1-150), proline-rich domain (PRD; aa:151-243), microtubule binding region (MTBR; aa:244-368) and C-terminus (aa:369-441). Scores are shown for *MAPT* KI controls, S305N, and P301S mice at 15M for the hippocampus (top panel, dataset#1) and the posterior cortex (bottom panel, dataset#2). Group differences reflect the relative burden of tau phosphorylation across genotypes. Data are presented as median and IQR (dashed grey line crosses the median of MAPT KI control). (**B**) Relative levels of phospho-tau peptides by targeted LC–MS analysis in the brains of 15M *MAPT* KI (N = 3), S305N (N = 3; ), and P301S (N = 3). Results are expressed in L/H ratio. (**C**) Representative images of coronal hemibrain sections used for the heatmap quantification in Fig. 3B from 6, 12, 15, 24 and 30M *MAPT* KI, S305N and P301S for AT8 IHC staining. Lower panels show zoomed-in boxes from 30M image from S305N and P301S mice.

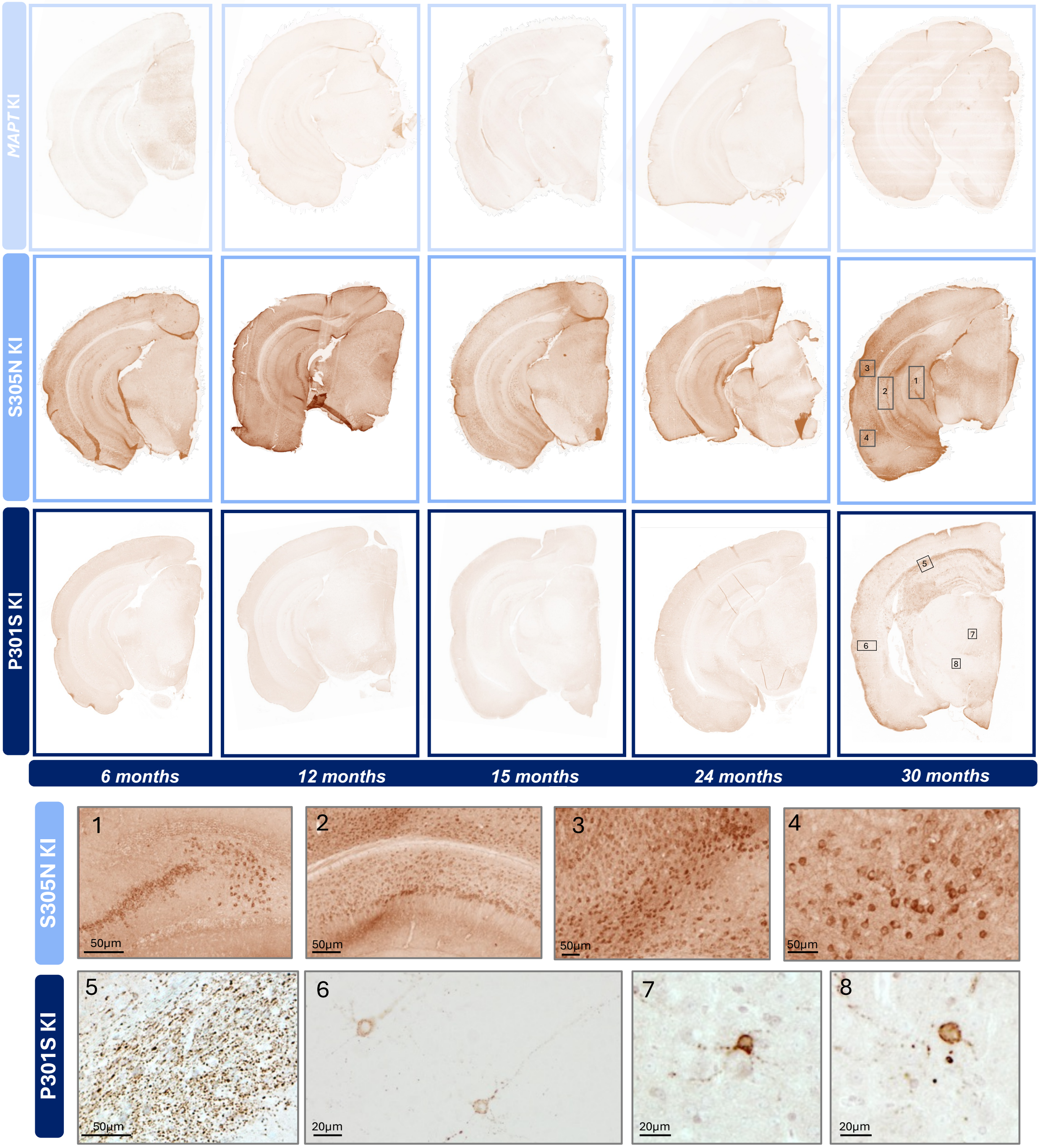

**Fig. S4.** Representative images of coronal hemibrain sections used for the heatmap quantification in **Fig. 4B** from 6, 12, 15, 24 and 30M *MAPT* KI, S305N and P301S for AT8 IHC staining. Lower panels show zoomed-in boxes from 30M image from S305N and P301S mice.

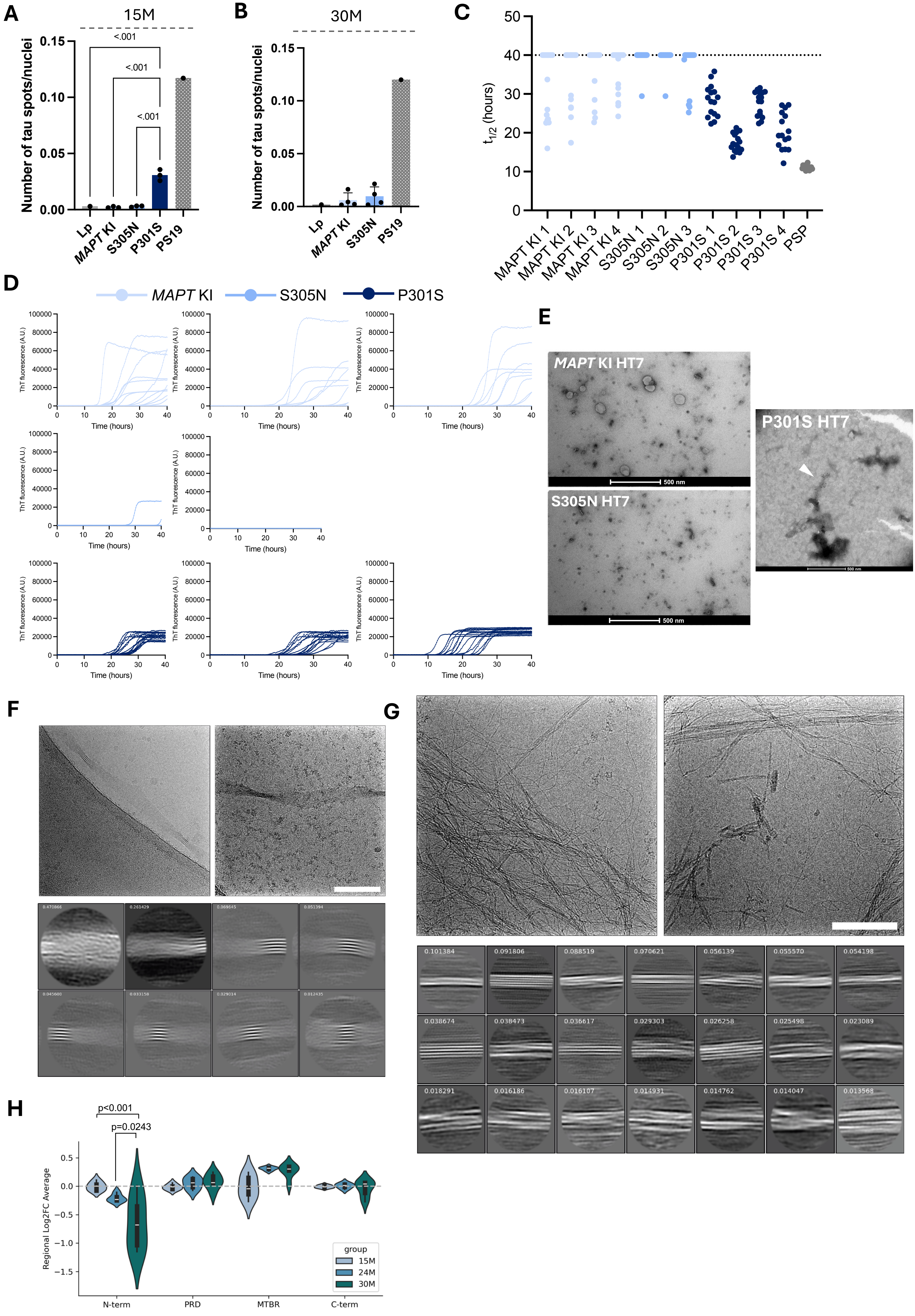
A
15M
Number of tau spots/nuclei
<.001
<.001
<.001
Lp
MAPT KI
S305N
P301S
PS19
B
30M
C
t1/2 (hours)
MAPT KI 1
MAPT KI 2
MAPT KI 3
MAPT KI 4
S305N 1
S305N 2
S305N 3
P301S 1
P301S 2
P301S 3
P301S 4
PSP
D
MAPT KI
S305N
P301S
ThT fluorescence (A.U.)
Time (hours)
E
MAPT KI HT7
S305N HT7
P301S HT7
500 nm
F
G
H
p<0.001
p=0.0243
Regional Log2FC Average
group
15M
24M
30M
N-term
PRD
MTBR
C-term

**Fig. S5. (A)** Tau seeding activity from 15M posterior cortex *MAPT* KI, S305N and P301S using S305N biosensor cells (N=3 per group; N=2 female and N=1 male), compared to PS19 mouse at 9M and lipofectamine negative control ("Lp"). Data represents mean±S.D. **(B)** Tau seeding activity from 15M posterior cortex *MAPT* KI and S305N using S305N biosensor cells (N=4 per group; sex-matched), compared to PS19 mouse at 9M and lipofectamine negative control ("Lp"). Data represents mean±S.D. **(C)** RT-QuIC aggregates half-way curve ($t_{1/2}$) from the combination of 16 replicates per biological sample from 15M posterior cortex from *MAPT* KI (N=4), S305N (N=3) and P301S (N=4; all sex-matched) and PSP positive control human brain (N=1). **(D)** Raw RT-QuIC ThT reactions from posterior cortex of mice at 15 months, *MAPT* KI (N=4), S305N (N=3) and P301S (N=4). N=1 is represented in Fig. 4E. **(E)** Representative negative-stain EM images of sarkosyl-insoluble tau fibrils isolated from whole-brain homogenates of 30M *MAPT* KI (top) and *S305N* mice (bottom). **(F)** Representative cryoEM micrographs and 2D class averages of sarkosyl-insoluble filaments (crossover distance 280-350 nm) purified from whole-brain P301S KI mouse sarkosyl-insoluble fraction at 30M. Scale bar represents 100 nm. **(G)** Representative cryoEM micrographs and 2D class averages of thin filaments (untwisted) purified from whole-brain P301S KI mouse sarkosyl-insoluble fraction at 30M. Scale bar represents 100nm. **(H)** Eigenprotein values for phospho-tau–associated peptides were derived from principal component analysis of TMT-based phospho-proteomics subdivided by tau regions, *i.e.* N-terminus (aa:1-150), proline-rich domain (PRD; aa:151-243), microtubule binding region (MTBR; aa:244-368) and C-terminus (aa:369-441). Scores are shown comparing hippocampal tissue from P301S mice at 15M, 24M and 30M. Group differences reflect the relative burden of tau phosphorylation across genotypes. Data are presented as median and IQR (dashed grey line crosses the median of *MAPT* KI control).

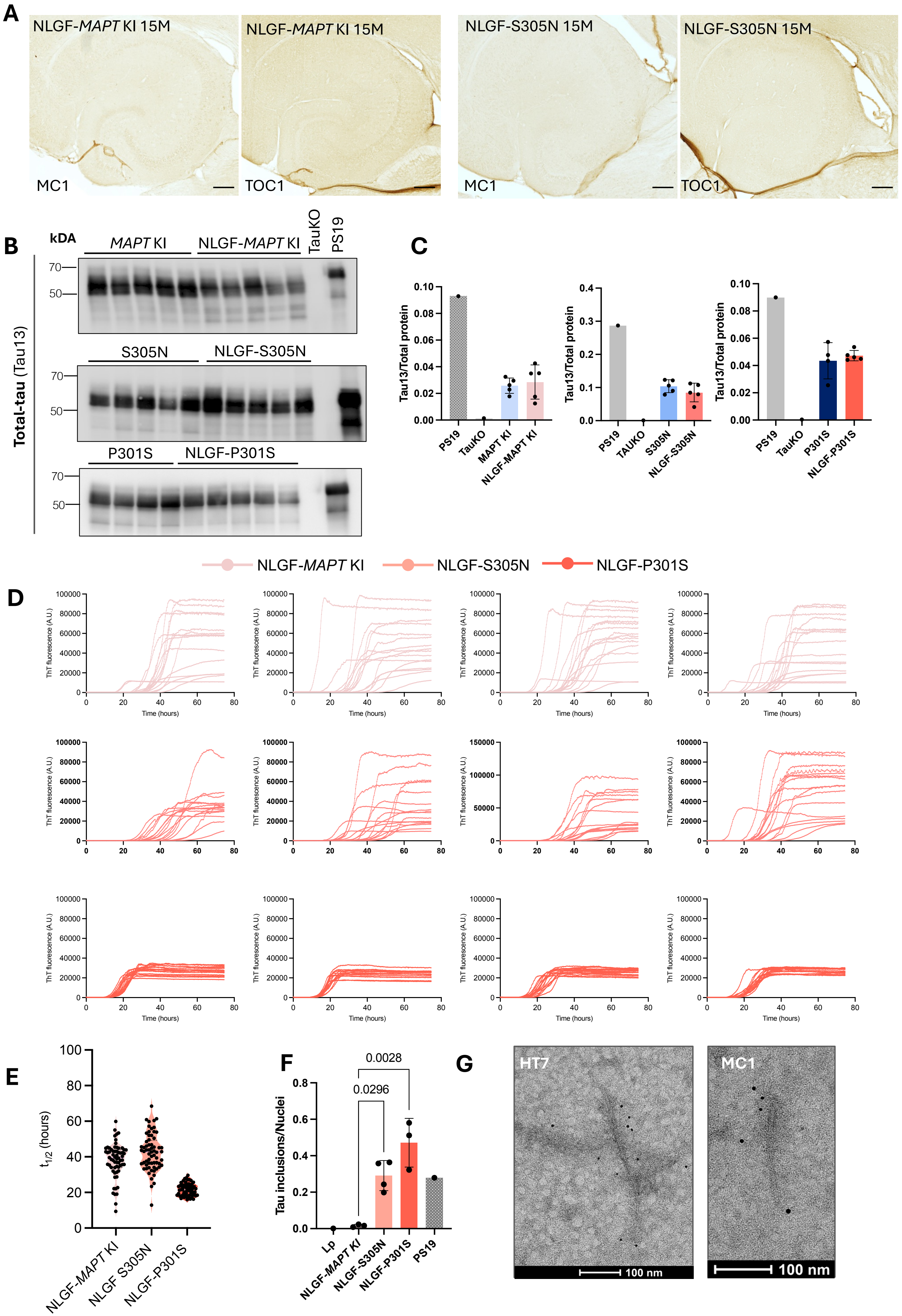

A
NLGF-MAPT KI 15M
MC1
NLGF-MAPT KI 15M
TOC1
NLGF-S305N 15M
MC1
NLGF-S305N 15M
TOC1
B
kDA
MAPT KI
NLGF-MAPT KI
TauKO
PS19
Total-tau (Tau13)
70
50
S305N
NLGF-S305N
P301S
NLGF-P301S
C
Tau13/Total protein
PS19
TauKO
MAPT KI
NLGF-MAPT KI
TAUKO
S305N
NLGF-S305N
P301S
NLGF-P301S
NLGF-MAPT KI
NLGF-S305N
NLGF-P301S
D
ThT fluorescence (A.U.)
Time (hours)
E
t1/2 (hours)
NLGF-MAPT KI
NLGF S305N
NLGF-P301S
F
Tau inclusions/Nuclei
0.0028
0.0296
Lp
NLGF-MAPT KI
NLGF-S305N
NLGF-P301S
PS19
G
HT7
MC1
100 nm
100 nm

**Fig. S6.** (**A**) Tau immunostaining detected by TOC1 (oligomeric tau) and MC1 (conformationally abnormal tau) antibodies in the hippocampus of NLGF-*MAPT* KI, NLGF-S305N at 15M. Scale bar represents 200µm. (**B**) Immunoblotting of total tau detected by Tau13 antibodies in the RIPA fraction of hippocampi brain lysates from NLGF-*MAPT* KI, NLGF-S305N and NLGF-P301S KI mice at 15M of age (N=5 for each group, expect for P301S with N=4; N=2-3 female and N=2 males), along with *Mapt* knock-out (KO) at 9M and positive control PS19 mice at 9M and (**C**) quantification to total protein. Data represents mean±S.D. (**D**) Raw RT-QuIC ThT reactions from posterior cortex of mice at 15M, NLGF-*MAPT* KI (N=4), NLGF-S305N (N=3) and NLGF-P301S (N=4; all sex-matched). (**E**) RT-QuIC aggregates half-way curve ($t_{1/2}$) from the combination of 16 replicates per biological sample from 15M posterior cortex from NLGF-*MAPT* KI (N=4), NLGF-S305N (N=3) and NLGF-P301S (N=4; all sex-matched). (**F**) Tau seeding activity from posterior cortex NLGF-*MAPT* KI, NLGF-S305N and NLGF-P301S (N=3 per group, N=2 female and N=1 male) using S305N biosensor cells nuclear inclusions in S305N biosensor at 30M, compared to PS19 mouse at 9M and lipofectamine negative control ("Lp"). Data represents mean±S.D. (**G**) Immunogold labelling (tau antibodies HT7 and MC1 on 6nm gold beads) of sarkosyl-insoluble extracted fibrils from whole-brain of 24M NLGF-P301S mice

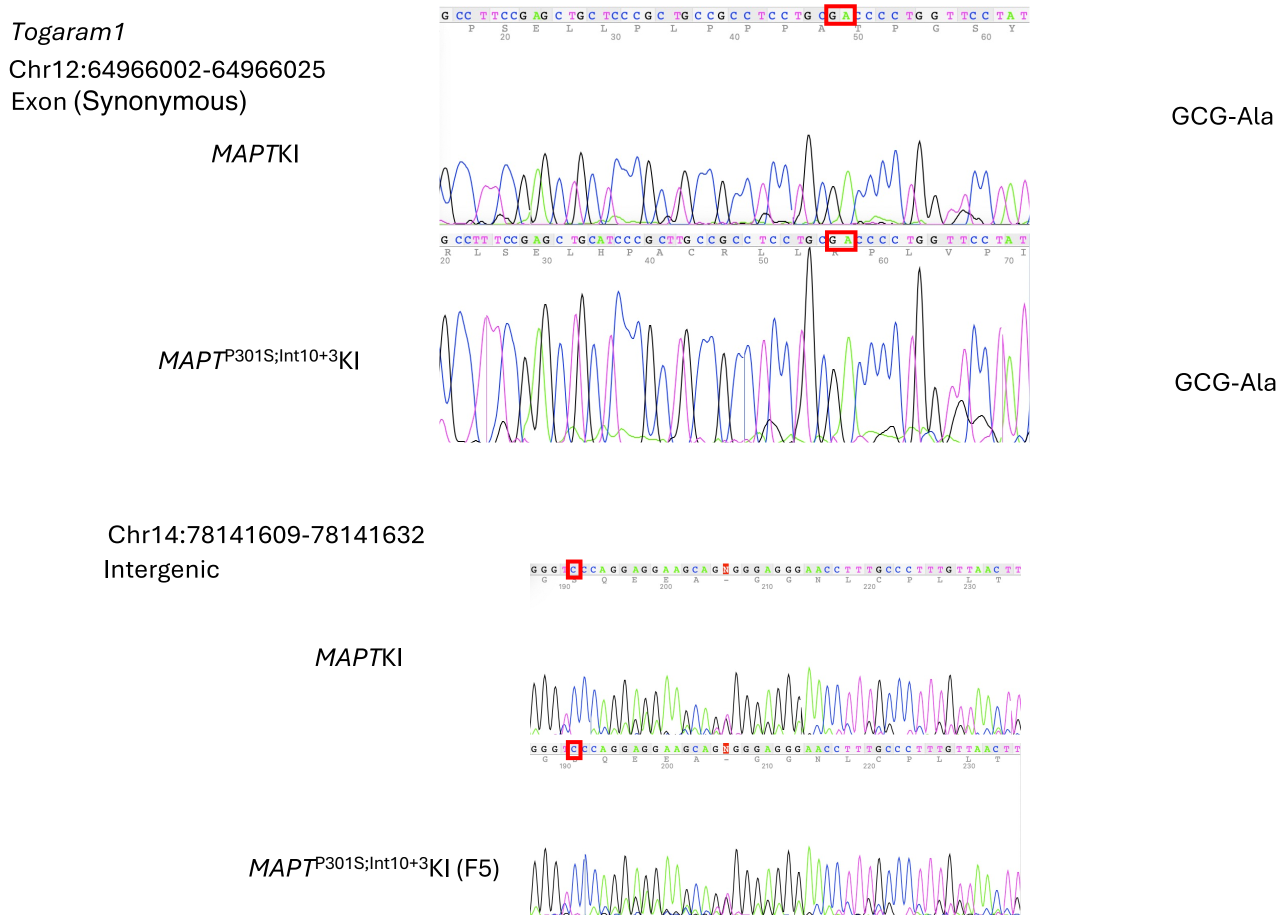

**Fig. S7** Off-target mutations identified in founders (highlighted in red boxes) were removed by backcrossing with C57B6/J mice more than four times.

**Table S1. Potential off-target sites for *MAPT* P301S;Intron10+3 line**

| Off-target candidate | Query type | Mismatch | Chr Position | Gene | Annotation | MAPT KIP301S; Inton10+3 G>A |
|---|---|---|---|---|---|---|
| **MAPT-P301-1** | No indel | 3 | Chr11:70351826-70351849 | Alox15 | Intron | - |
| **MAPT-P301-2** | No indel | 3 | Chr4:119200276-119200299 | Svbp | Intron | - |
| **MAPT-P301-3** | No indel | 3 | Chr9:64060473-64060496 | | Intergenic | - |
| **MAPT-P301-4** | No indel | 3 | Chr6:95725509-95725532 | | Intergenic | - |
| **MAPT-P301-5** | No indel | 3 | Chr12:64966002-64966025 | Togaram1 | Exon | ○ |
| **MAPT-P301-6** | No indel | 3 | Chr14:78141609-78141632 | | Intergenic | ○ |
| **MAPT-P301-7** | No indel | 3 | Chr2:126701111-126701134 | | Intergenic | - |
| **MAPT-P301-8** | No indel | 3 | Chr12:72964402-72964425 | | Intergenic | - |
| **MAPT-P301-9** | No indel | 3 | Chr1:180813179-180813202 | H3f3a | Intron | - |
| **MAPT-P301-10** | No indel | 3 | Chr6:146888630-146888653 | Ppfibp1 | Non-coding | - |
| **MAPT-P301-11** | Del 8 | 2 | Chr9:121021200-121021222 | Ulk4 | Intron | - |
| **MAPT-P301-12** | Del 3 | 2 | ChrX:87776462-87776484 | Il1rapl1 | Intron | - |
| **MAPT-P301-13** | Del 2 | 2 | Chr16:18687059-18687081 | | Intergenic | - |
| **MAPT-P301-14** | Del PAM 2 | 2 | Chr2:167736542-167736564 | | Intergenic | - |
| **MAPT-P301-15** | Ins 17 | 2 | Chr10:118675703-118675727 | | Intergenic | - |
| **MAPT-P301-16** | Ins 16 | 2 | Chr2:133528883-133528907 | | Intergenic | - |
| **MAPT-P301-17** | Ins 13 | 2 | Chr14:26429735-26429759 | Slmap | Intron | - |
| **MAPT-P301-18** | Ins 7 | 2 | Chr6:146527101-146527125 | | Intergenic | - |
| **MAPT-P301-19** | Ins 5 | 2 | Chr2:131262710-131262734 | Pank2 | Intron | - |
| **MAPT-intron10+3 G>A-1** | No indel | 3 | Chr14:31275639-31275661 | Dnah1 | Intron | - |
| **MAPT-intron10+3 G>A-2** | No indel | 3 | Chr10:67285234-67285256 | Nrbf2 | Non-coding | - |
| **MAPT-intron10+3 G>A-3** | No indel | 3 | Chr3:108148064-108148086 | | Intergenic | - |
| **MAPT-intron10+3 G>A-4** | No indel | 3 | Chr5:137062275-137062297 | Serpine1 | Non-coding | - |
| **MAPT-intron10+3 G>A-5** | No indel | 3 | Chr6:72577777-72577799 | Elmod3 | Intron | - |
| **MAPT-intron10+3 G>A-6** | No indel | 3 | Chr5:64508703-64508725 | | Intergenic | - |
| **MAPT-intron10+3 G>A-7** | Del 18 | 2 | Chr6:59413774-59413795 | Gprin3 | Intron | - |

| | | | | | | |
|---|---|---|---|---|---|---|
| **MAPT-intron10+3 G>A-8** | Del 18 | 2 | Chr7:96207591-96207612 | Tenm4 | Intron | - |
| **MAPT-intron10+3 G>A-9** | Del 17 | 2 | Chr2:168700314-168700335 | Atp9a | Intron | - |
| **MAPT-intron10+3 G>A-10** | Del 16 | 2 | Chr2:146621362-146621383 | | Intergenic | - |
| **MAPT-intron10+3 G>A-11** | Del 16 | 2 | Chr8:69915135-69915156 | Gatad2a | Intron | - |
| **MAPT-intron10+3 G>A-12** | Del 16 | 2 | Chr11:99056068-99056089 | | Intergenic | - |
| **MAPT-intron10+3 G>A-13** | Del 15 | 2 | Chr2:167736538-167736559 | | Intergenic | - |
| **MAPT-intron10+3 G>A-14** | Del 14 | 2 | Chr1:82288807-82288828 | Irs1 | Exon | - |
| **MAPT-intron10+3 G>A-15** | Del 14 | 2 | Chr4:150626514-150626535 | | Intergenic | - |
| **MAPT-intron10+3 G>A-16** | Del 14 | 2 | Chr17:30516288-30516309 | Btbd9 | Intron | - |
| **MAPT-intron10+3 G>A-17** | Del 14 | 2 | Chr8:112093736-112093757 | | Intergenic | - |
| **MAPT-intron10+3 G>A-18** | Del 14 | 2 | Chr12:84648519-84648540 | Vrtn | Exon | - |
| **MAPT-intron10+3 G>A-19** | Del 14 | 2 | Chr11:87664708-87664729 | Rnf43 | Exon | - |
| **MAPT-intron10+3 G>A-20** | Del 13 | 2 | Chr4:139161305-139161326 | Gm33304 | Non-coding | - |
| **MAPT-intron10+3 G>A-21** | Del 12 | 2 | Chr18:80778335-80778356 | Atp9b | Intron | - |
| **MAPT-intron10+3 G>A-22** | Del 12 | 2 | Chr2:119770882-119770903 | Rpap1 | Intron | - |
| **MAPT-intron10+3 G>A-23** | Del 12 | 2 | Chr6:109857978-109857999 | | Intergenic | - |
| **MAPT-intron10+3 G>A-24** | Del 12 | 2 | Chr5:142468117-142468138 | Ap5z1 | Exon | - |
| **MAPT-intron10+3 G>A-25** | Del 11 | 2 | Chr17:25386575-25386596 | Cacna1h | Intron | - |
| **MAPT-intron10+3 G>A-26** | Del 11 | 2 | Chr9:119555524-119555545 | Scn5a | Intron | - |
| **MAPT-intron10+3 G>A-27** | Del 9 | 2 | Chr9:43536132-43536153 | | Intergenic | - |
| **MAPT-intron10+3 G>A-28** | Del 9 | 2 | Chr7:125974764-125974785 | Gsg1l | Intron | - |

| | | | | | | |
|---|---|---|---|---|---|---|
| **MAPT-intron10+3 G>A-29** | Del 6 | 2 | Chr12:91965237-91965258 | | Intergenic | - |
| **MAPT-intron10+3 G>A-30** | Del 4, or Del 5 | 2 | Chr11:118367373-118367394 | | Intergenic | - |
| **MAPT-intron10+3 G>A-31** | Del 4, or Del 5 | 2 | Chr4:43394834-43394855 | Rusc2 | Intron | - |
| **MAPT-intron10+3 G>A-32** | Del 4, or Del 5 | 2 | Chr5:121818513-121818534 | Sh2b3 | Exon | - |
| **MAPT-intron10+3 G>A-33** | Del 3 | 2 | Chr12:21295846-21295867 | Cpsf3 | Intron | - |
| **MAPT-intron10+3 G>A-34** | Del 3 | 2 | Chr9:113870747-113870768 | Clasp2 | Intron | - |
| **MAPT-intron10+3 G>A-35** | Del 3 | 2 | Chr6:125633288-125633309 | Vwf | Intron | - |
| **MAPT-intron10+3 G>A-36** | Del 3 | 2 | Chr2:130840193-130840214 | 4930402H24Rik | Non-coding | - |
| **MAPT-intron10+3 G>A-37** | Del 1, or Del 2 | 2 | Chr17:49049099-49049120 | Lrfn2 | Intron | - |
| **MAPT-intron10+3 G>A-38** | Ins 17 | 2 | Chr16:22558979-22559002 | Dgkg | Intron | - |
| **MAPT-intron10+3 G>A-39** | Ins 16 | 2 | Chr15:74631843-74631866 | Mroh4 | Intron | - |
| **MAPT-intron10+3 G>A-40** | Ins 15 | 2 | Chr17:54077434-54077457 | | Intergenic | - |
| **MAPT-intron10+3 G>A-41** | Ins 13 | 2 | Chr7:139648287-139648310 | Cfap46 | Intron | - |
| **MAPT-intron10+3 G>A-42** | Ins 13 | 2 | Chr2:30194586-30194609 | Kyat1 | Intron | - |
| **MAPT-intron10+3 G>A-43** | Ins 6 | 2 | Chr8:71290281-71290304 | Myo9b | Non-coding | - |
| **MAPT-intron10+3 G>A-44** | Ins 6 | 2 | Chr5:138859529-138859552 | | Intergenic | - |
| **MAPT-intron10+3 G>A-45** | Ins 6 | 2 | Chr16:93903368-93903391 | Chaf1b | Intron | - |
| **MAPT-intron10+3 G>A-46** | Ins 1 | 2 | Chr13:63111073-63111096 | Aopep | Intron | - |

**Table S2. Primer sequences for whole-genome resequencing.**

| *MAPT*-P301-5 |
|---|

Togaram1 F AATGGTTCCCTCCAGCTCTC
Togaram1 R TTTTTCTCGCCTCTCATGGT

| *MAPT*-P301-6 |
|---|

Intergenic p301s
F AGTGTCCTTTGTCATGACCACA
Intergenic p301s
R GGGACTTACATGTTTCACTCAGG

**Table S3. Antibody list**

The following antibodies were used at the dilutions described below.

| | Antibodies | Source | WB (dilution) | IHC (dilution) | Other (dilution) |
|---|---|---|---|---|---|
| **Primary** | CP13 | Kindly provided by Cristina D'Abramo | 1:500 | 1:500 | |
| | AT8 | Thermo Fisher Scientific (MN1020B) | 1:500 | 1:500 | |
| | TOC1 | Kindly provided by Nicholas Kanaan | 1:5000 | 1:1000 | |
| | MC1 | Kindly provided by Cristina D'Abramo | 1:500 | 1:500 | Immunogold labelling 1:2 |
| | PHF1 | Kindly provided by Cristina D'Abramo | | 1:500 | |
| | Tau13 | Santa Cruz (sc-21796) | 1:2000 | | |
| | Tau5 | Thermo Fisher Scientific (AHB0042) | 1:2000 | | |
| | HT7 | Themo Fischer Scientific (MN1000) | | | Immunogold labelling 1:2 |
| | Synaptotagmin | Synaptic System (105-002) | | 1:500 | |
| | Homer 1 | Synaptic System (160-004) | | 1:500 | |
| **Secondary** | Goat anti-chicken 488 Alexa Fluor™ | Thermo Fisher Scientific (A-11039) | | 1:500 | |
| | Goat anti-rabbit 594 Alexa Fluor™ | Thermo Fisher Scientific (A-11012) | | 1:500 | |
| **Stains** | Hoechst 33342 | Thermo Fisher Scientific | | 1:5000 | |
| | Gallyas Silver stain | | | | |

**Table S4. Real-time PCR for primers**

3R-Tau F GTCCGTACTCCACCCAAGTC
3R-Tau R TTTGTAGACTATTTGCACCTTCC
4R-Tau F GAAGCTGGATCTTAGCAACG
4R-Tau R GACGTGTTTGATATTATCCT
Total Tau F AGCCAAGACATCCACACGTT
Total Tau R ATCAGAGGGTCTGAGCTACCA
G3PDH F CCATGGCACCGTCAAGGCTGA
G3PDH R GCCAGTAGAGGCAGGGATGAT

**Table S5. Custom primers used for RNA extraction and PCR**

Fwd: AAGTCGCCGTCTTCCGCCAAG

Rev: GTCCAGGGACCCAATCTTCGA

**Table S6**

| Cluster ID | ABI2023_subclass | Custom_name | Region | MainNT | subclass_color |
|---|---|---|---|---|---|
| 0 | DG Glut | DG | HPF | Glut | #CC2400 |
| 1 | L2/3 IT CTX Glut | L2/3 IT CTX | CTX | Glut | #E9530F |
| 2 | L4/5 IT CTX Glut | L4/5 IT CTX | CTX | Glut | #3DCCB7 |
| 3 | CA1-ProS Glut | CA1-ProS | HPF | Glut | #0F3466 |
| 4 | L6 CT CTX Glut | L6 CT CTX | CTX | Glut | #34661F |
| 5 | PVALB Gaba | Pvalb | MGE | GABA | #661F3B |
| 6 | CA3 Glut | CA3 | HPF | Glut | #C973FF |
| 7 | SST Gaba | Sst | MGE | GABA | #99F2FF |
| 8 | MEA Slc17a7 Glut,COAp Grxcr2 Glut,LA-BLA-BMA-PA Glut | Amyg | Amygdala | Glut | #456C99 |
| 10 | VIP Gaba, Sncg Gaba, RHP-COA Ndnf Gaba | Vip/Sncg | CGE | GABA | #663D47 |
| 11 | L6b CT ENT Glut,L6b CT CTX Glut | L6b CT | CTX/ENT | Glut | #9EFF99 |
| 12 | L6 IT CTX Glut | L6 IT CTX | CTX | Glut | #C0FF4D |
| 13 | L5 IT CTX Glut | L5 IT CTX | CTX | Glut | #660F38 |
| 14 | L2/3 IT RSP Glut | L2/3 IT RSP | RSP | Glut | #999145 |
| 15 | L2/3 IT PPP Glut | L2/3 IT PPP | PPP | Glut | #925C99 |
| 16 | L2/3 IT ENT Glut | L2/3 IT ENT | ENT | Glut | #1F46CC |
| 17 | L5 NP Ctx Glut | L5 NP CTX | CTX | Glut | #7B7ACC |
| 18 | LSX Nkx2-1 Gaba,LSX Prdm12 Slit2 Gaba, LSX Prdm12 Zeb2 Gaba, 072 LSX Sall3 Lmo1 Gaba | LSX Gaba | Lateral Septal Complex | GABA | #3283FE |
| 19 | SUB-ProS Glut | SUB-ProS | HPF | Glut | #FF9999 |
| 21 | L5 ET CTX Glut | L5 ET CTX | CTX | Glut | #99A2FF |
| 22 | CA3 Glut | CA3 | HPF | Glut | #C973FF |
| 23 | STR D1 Gaba,STR D2 Gaba | STR Gaba | Striatum | GABA | #005B66 |
| 24 | Lamp5 Gaba | Lamp5 | MGE | GABA | #FF764D |
| 25 | CT SUB Glut | CT SUB | HPF | Glut | #440066 |
| 26 | L2/3 IT PIR-ENTl Glut,L2 IT ENT-po Glut | L2/3 IT PIR/ENTl-po | ENT | Glut | #940099 |
| 27 | L2/3 IT PPP Glut | L2/3 IT PPP | PPP | Glut | #925C99 |
| 28 | L4 RSP-ACA Glut | L4 RSP-ACA | RSP-ACA | Glut | #FFF226 |
| 29 | Lamp5 Lhx6 Gaba, Pvalb Chandelier Gaba | Lamp5 Lhx6 / Pvalb Chandelier | MGE/CGE | GABA | #2F992E |
| 31 | ENTmv-PA-COAp Glut | ENTmv-PA-COAp | ENT | Glut | #B6CC7A |
| 32 | CLA-EPd-CTX Car3 Glut | CLA-EPd-CTX Car3 | | Glut | #64c2fc |
| 33 | L2/3 IT PPP Glut | L2/3 IT PPP | PPP | Glut | #925C99 |
| 34 | HPF CR Glut | HPF CR | HPF | Glut | #919900 |

| | | | | | |
|---|---|---|---|---|---|
| **35** | L2 IT PPP-APr Glut | L2 IT PPP-APr | PPP | Glut | #0F6632 |
| **36** | STR-PAL Chst9 Gaba | STR-PAL Chst9 | Striatum | GABA | #967B09 |
| **39** | L5 ET CTX Glut | L5 ET CTX | CTX | Glut | #99A2FF |
